# Highly resolved tumor architecture via matched spatial and nucleus transcriptomics from a single tissue section

**DOI:** 10.64898/2026.09.07.749790

**Authors:** Marcos Tito Machado, Mengxiao He, Leire Alonso Galicia, Zaneta Andrusivova, Emmanouela Perisynaki, Matthew W Myers, Sarantis Giatrellis, Sandra O’Toole, Beata Kiedik, Raphaël Mauron, Sophie van der Leij, Kate Harvey, John Reeves, Javier Escudero Morlanes, Taobo Hu, Mengping Long, Mats Nilsson, Tianyi Li, Xinsong Chen, Johan Hartman, Máté Mihálffy, Taopeng Wang, Marco Vicari, Laura Savolainen, Andrew Erickson, Sandy Figiel, Alastair Lamb, Alexander Swarbrick, Joakim Lundeberg, Reza Mirzazadeh

**Author notes:** These authors contributed equally to this work (Marcos Tito Machado, Mengxiao He, Leire Alonso Galicia).

## Abstract

Spatial transcriptomics often relies on reference-based deconvolution to infer cell types in a tissue context; however, public single-cell datasets can miss patient-specific biology. Here we introduce SIMPlex, a method that generates matched spatial and single-nucleus gene-expression profiles from the same 5 µm FFPE section. We demonstrate context-matched profiles across mouse brain, breast cancer and prostate cancer tissues, resolving fine-grained cell-states with distinct spatial signatures. By extracting both spatial and nuclear layers, SIMPlex maximises the information recovered from a single tissue section, an advantage for scarce archival and clinical specimens.

## Main Text

Sequencing-based spatial transcriptomic assays have transformed tissue profiling by mapping gene expression in histological context. However, due to inherent cell mixing and signal sparsity at the spot/bin level, computational deconvolution or imputation is typically required to infer cell type contribution^1^. These generally require a single-cell/nucleus RNA sequencing (sc/snRNA-seq) reference, commonly obtained from external public atlases^2,3^. These may not capture the biology of the profiled section, particularly in heterogeneous tumors, where differences in platform, preservation, and patient context add further discordance^4–7^. Generating a matched snRNA reference is especially challenging for routine clinical Formalin-Fixed Paraffin-Embedded (FFPE) material. Clinical archives retain FFPE blocks and slides, but tissue block access for research is often limited, making direct snRNA-seq from thin sections difficult^8,9^.

To address this need, we developed SIMPlex (Single-section Integrated Multilayer Profiling), an experimental workflow that enables recovery of nuclei from post-Visium FFPE tissue sections for matched-cell-state analysis (via emulsion snRNA-seq). The protocol is demonstrated on routine clinical 5 µm FFPE sections. Same-section multimodal profiling has previously been demonstrated in fresh-frozen and FFPE tissue^10–14^, and scRNA-seq based deconvolution is well established. Recently, patient-matched protocols have been developed to further refine the mapping of cell types in spatial transcriptomics data^7^; however, these require large samples or consecutive tissue scrolls to generate sufficient data and are therefore less attractive for limited and archival material.

After the spatial workflow, the sections are scraped and hybridized with transcriptome-wide probes, and nuclei are recovered and processed by standard droplet-based microfluidics. The method requires no modifications to the spatial protocol, supports long-term slide storage prior to nuclei recovery, and operates across both standard and high-resolution spatial assays as well as fresh-frozen (FF) tissues. (**Fig. 1a; Extended Data Fig. 1a**). We validated the method using 5 µm FFPE breast tumors (n = 8) and prostate tumors (n = 2), alongside 18 µm FF breast tumors and 12 µm FF mouse brain (**Fig. 1b**), establishing its generalizability across tissues, preservation methods, and section thicknesses.

**Fig. 1.**
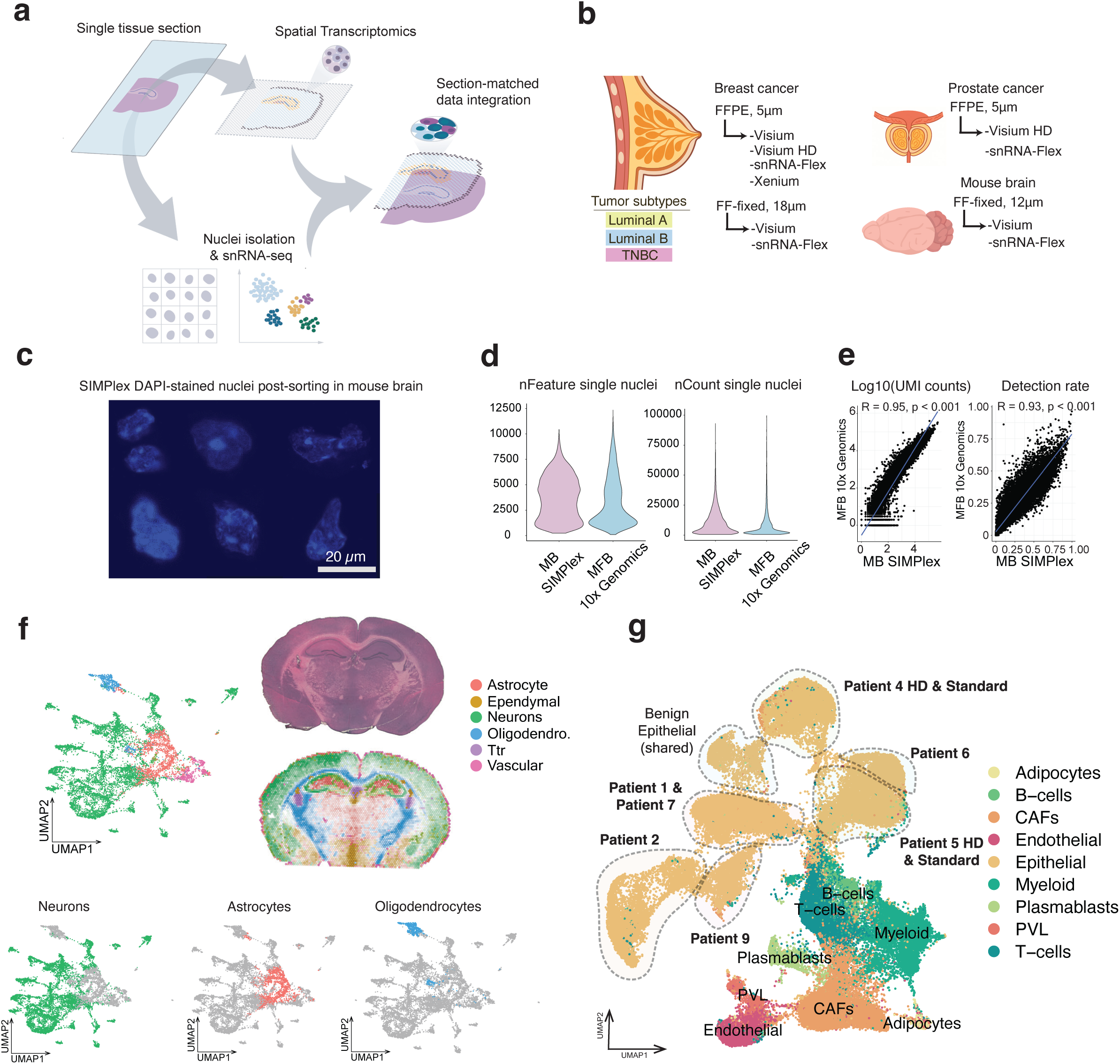
SIMPlex and validation across tissues. **a,** Workflow: Spatial transcriptomics of a single H&E section, followed by sorted nuclei isolation for matched snRNA-seq and subsequent matched multimodal data integration. **b,** Overview of datasets and profiling platforms. **c,** Representative SIMPlex DAPI-positive nuclei from sorted mouse brain sections. **d,** Distributions of unique genes (nFeature) and molecules (nCount) per nucleus visualized as violin plots; MB (mouse brain), MFB (mouse forebrain) **e,** Gene-gene scatter plot of log10 transformed UMI counts and detection rate plot. The detection rate for a gene is defined as the proportion of spots with detected UMI counts; MB (mouse brain), MFB (mouse forebrain). **f,** UMAP of the SIMPlex snRNA-seq data and deconvolution of the matched section. **g,** Integrated UMAP of all SIMPlex nuclei from 8 BC patients, colored by major cell types. Dashed outlines denote patient-specific tumor and benign epithelial populations.

We first benchmarked SIMPlex in the mouse brain, a tissue with well-defined spatial organisation. After spatial transcriptomics, we recovered 11,129 barcoded nuclei from a 12 μm FF section which resulted in quality metrics comparable to manufacturer datasets, with a median of 2,100 detected genes and 3,500 UMIs per nucleus and strong gene-level concordance (R = 0.95 for UMI counts; R = 0.93 for detection rate), demonstrating preserved nuclear RNA (**Fig. 1c-e; Extended Data Fig. 1b-d**). Label transfer from public references^15,16^, confirmed expected neuronal, oligodendrocyte, and astrocyte domains **(Fig. 1f; Extended Data Fig. 1e**), and recovered layer-specific neuronal subtypes and cortical and hippocampal lamination (**Extended Data Fig. 1f-g**). Together, these results demonstrate that nuclei recovered after spatial transcriptomics retain sufficient integrity for accurate mouse brain cell-type mapping.

Next, we extended the application to both FF and FFPE human breast cancer (BC) tissue. In FF samples, we first assessed nuclei profiles by sorting and found that DAPI-positive intact nuclei distribution decreased with tissue thickness (**Extended Data Fig. 2a**). We then performed SIMPlex on two 18 µm FF sections, both of which yielded robust data (**Extended Data Fig. 2b-e**). Clinical routine 5 µm sections obtained from archival FFPE tumor blocks yielded median 7678 barcoded nuclei per section after sequencing and QC filtering (**Supplementary Table 1**). Despite post-Visium run and presence of partial nuclei, median values of 950 UMIs and 789 genes per nucleus were comparable to published FFPE sc/snRNA-seq datasets^7^ (**Extended Data Fig. 2f-h, Extended Data Fig. 3**). FFPE prostate tumors showed similar performance (**Extended Data Fig. 4**), indicating that standard pathology thickness of typically large tissue sections is adequate for generating snRNA-seq across tissues. The applicability of the protocol to other clinical samples, such as biopsies remains to be determined, albeit stacking of needle biopsies to obtain a proxy of a larger tissue can be foreseen^17^.

Having established technical feasibility over multiple tissue types, we next assessed the biological granularity achieved by our method in FFPE BC tissue. Across eight tumor samples spanning triple-negative (TNBC), luminal A, and luminal B subtypes, SIMPlex generated a dataset of 71,418 nuclei resolving all major epithelial, immune, and stromal lineages (**Fig. 1g; Extended Data Fig. 5a**). These profiles recapitulated key populations reported in breast cancer atlases^2,3^. Malignant epithelial states segregated by patient and subtype, while stromal, immune, and benign epithelial compartments were more conserved across patients. Furthermore, same-patient consecutive sections (patients 4 and 5) integrate seamlessly prior to batch-correction, suggesting high reproducibility, enabling coherent interpretation of these states (**Extended Data Fig. 5a-c**). Comparison of histological compartments across BC samples demonstrates better cell type concordance, as expected of section-matched data, when compared to the bulk-dissociated single-cell BC Atlas reference^3^ (**Fig. 2a**).

**Fig. 2.**
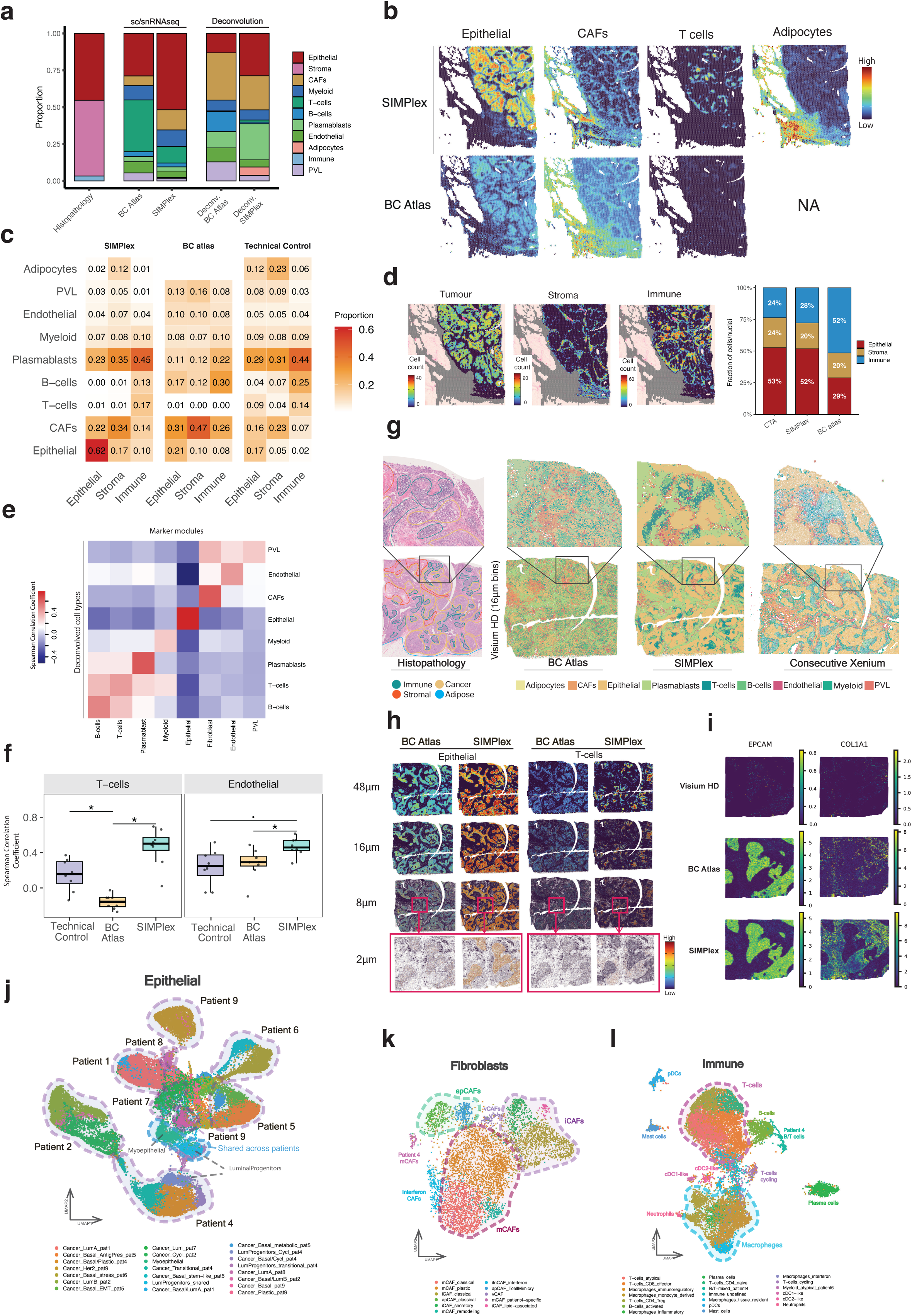
Mapping of BC tumor microenvironment and exploration of high-resolution cell states. **a,** Bulk overview population composition for histopathology, single-cell/nuclei labels, and spatial deconvolution. Each color represents a cell type or histopathology annotation compartment. **b,** Spatial visualization of deconvolution proportions for major cell types. **c,** Cell type proportions within main histopathological labels using different references. **d,** Spatial visualization of CTA assignments displayed as cell counts for main populations in the tumor microenvironment, and tissue compartment composition (fraction of cells/nuclei) for the CTA ground truth, SIMPlex, and the BC atlas. **e,f,** Benchmarking SIMPlex deconvolution accuracy against UCell gene signature scoring (**e)** between deconvolved cell type abundances (using different references) and marker gene scores. (**f**) T cell and Endothelial correlation coefficients across patients (Wilcoxon signed-rank test and Benjamini-Hochberg correction; *: *FDR* < 0.05; ●: *FDR* < 0.1). **g,h,** Benchmarking SIMPlex performance on high-resolution mapping. Spatial maps of (**g**) dominant cell types mapped to 16 μm bins using SIMPlex and BC Atlas, compared with a consecutive Xenium section with colour-matched labels (cell types obtained via label transfer from BC Atlas). **(h)** major cell types using different references across resolution bins. **(i)** Spatial gene imputation of EPCAM and COL1A1 using section-matched SIMPlex data and BC atlas as references at 8um resolution. **j-l,** Subclustering of BC populations (**j)** Epithelial populations colored by cluster and dashed outlines represent patient-specific or shared identities. (**k)** Fibroblasts with dashed outlines representing main and patient-specific CAF phenotypes. (**l**) Immune populations with dashed outlines reflecting main and patient-specific cell types.

Consistent with histopathology, section-matched cell type deconvolution accurately reflected expected tissue architecture, mapping adipocytes and cancer-associated fibroblasts (CAFs) to stromal regions, epithelial cells to tumor compartments, and T cells to focal immune aggregates. In contrast, BC Atlas produced less precise cell type mappings as exemplified at tumor-stroma-immune boundaries, including the CAF signal mixing into adipocyte-rich regions. Notably, the BC atlas lacked adipocyte mapping entirely due to their absence in the reference, a common limitation of scRNA-seq references due to dissociation challenges **(Fig. 2b,c; Extended Data Fig. 5d and 6a)**. In addition, snRNA-seq on non-Visium adjacent tissue sections from patients 4 and 6 failed to recover sufficient genes per nucleus for confident downstream analysis (**Extended Data Fig. 2h**). To further assess the impact of patient and section-specific matching on deconvolution, we compared three reference strategies: section-matched, the public scRNA-seq BC Atlas^3^, and a Technical Control reference. The latter aggregated nuclei from all other patients, excluding the matched case, to isolate the effect of patient-specific biology while preserving identical probe chemistry. We benchmarked deconvolution performance across those three references using known marker genes and computational tissue annotation^18^ (CTA). CTA provided an orthogonal image-based validation, where our matched data achieved superior spot-level concordance relative to other references, as well as closely recapitulating compartment-level proportions (**Fig. 2d**; **Extended Data Fig. 6b-d**). When compared to known cell type marker genes, deconvolution with section-matched, showed an overall increase in correlation; however, only T cells and endothelial reach statistical significance (*FDR < 0.05*) compared to the reference atlas. For most cell types, given the major annotation resolution, it is expected that both SIMPlex and Technical Control data recapitulate BC Atlas spatial distribution **(Fig. 2e,f; Extended Data Fig. 7a-d).**

To evaluate performance at near single-cell resolution, we applied our developed methodology to BC sections profiled with Visium HD and performed Xenium in situ profiling on consecutive sections as an independent validation. With section-matched nuclei references, deconvolution of 16 µm bins preserved tumor architecture and stromal-immune interfaces, aligning closely with histology and in situ profiling, whereas atlas-based references produced less precise patterns (**Fig. 2g; Extended Data Fig. 7e**). Across different resolution bins (48 µm - 2 µm), SIMPlex sharpened epithelial and immune boundaries and maintained focal lymphoid niches (**Fig. 2h; Extended Data Fig. 7f**). Visium HD data is notably sparse at high resolution bins. SIMPlex data can be leveraged for gene expression imputation at 8 µm bin resolution, demonstrating its value for recovering contextual biological signatures on poor/sparse spatial data. SIMPlex reconstructions recapitulate BC atlas on markers for most lineages, and data suggests that it could provide improvements for CAF expression (*COL1A1*) (**Fig. 2i**). Together, these results show high-fidelity mapping and reconstruction using sparse Visium HD data across different resolutions.

Having validated the methodology at the major cell type level, we next resolved finer, context-specific transcriptional states. Subclustering epithelial, immune, and fibroblast clusters from >60,000 nuclei, we identified 55 transcriptionally distinct subpopulations, generating a high-resolution collection of heterogeneous inter-patient cell states, 23 of which were patient-specific. (**Fig. 2j-l; Extended Data Fig. 8a-c**). SIMPlex demonstrates clear patient-specific epithelial clusters, consistent with previous studies^2,3^, while resolving transcriptionally distinct tumor-associated expression programs. These patterns reflect biological diversity spanning subtype-restricted, transitional, and shared normal epithelial states such as myoepithelial cells and luminal progenitors (**Fig. 2j**; **Extended Data Fig. 8d)**. Stromal analysis revealed matrix-remodeling, inflammatory, and antigen-presenting CAF (apCAF) phenotypes^19–22^, alongside interferon-related and a patient 4-specific CAF state (**Fig. 2k**; **Extended Data Fig. 8e**). Immune profiling showed macrophage and T cell-dominated landscapes with regulatory and interferon-active phenotypes, consistent with spatially organised immune niches in BC^2,3,23^ (**Fig. 2l; Extended Data Fig. 8f**). Notably, SIMPlex resolved mast cell and neutrophil populations, rarely captured in other atlases^2,3^ due to possible dissociation challenges associated with single-cell^24^ (**Fig. 2l).** Together, these data show that SIMPlex preserves fine-grained epithelial, stromal, and immune heterogeneity shaped by each tumor microenvironment, capturing context-dependent states that are likely attenuated in conventional scRNA-seq references.

To assess whether transcriptional specificity is preserved spatially, we mapped SIMPlex-defined cell states using data pooled from all single-nuclei. Section-matched mapping confirmed that patient-specific tumor programs localized almost exclusively to their tissue of origin **(Fig. 3a)**. We then further examined tumor-microenvironment structure in Patient 4, a TNBC case containing both Ductal carcinoma in situ (DCIS) and invasive carcinoma, using section-matched deconvolution, cell state colocalization, and inferred spatial niches^25^ **(Fig. 3b,c; Extended Data Fig. 9a-c)**. Section-matched data resolved distinct cell states into niches with specific epithelial-stromal co-localizations that aligned with histopathological annotations **(Fig. 3c,d; Extended Data Fig. 9d)**. Cell state abundances were visualized as a function of distance from the DCIS/Invasive border, revealing boundary-associated gradients and enrichments such as DCIS-adjacent plasmacytoid dendritic cells (pDCs) and apCAFs surrounding the invasive core **(Fig. 3e)**. Besides displaying established mCAF-tumor and iCAF-stromal regionalizations^22,26^, our data resolved heterogeneity within each of these phenotypes, with mCAF segregating into distinct spatial profiles: a DCIS-confined remodeling profile, and invasive-associated states (**Fig. 3f; Extended data Fig. 9e**). iCAF segmented into tumor-bordering iCAFs, close to immune-rich niches, and secretory iCAFs located in the non-invasive stroma **(Fig. 3g)**. The BC atlas did not recapitulate this degree of spatial segregation at the finer population level **(Extended Data Fig. 9f)**, and differential expression analyses confirm gene expression signatures of patient-specific cell states **(Extended Data Fig. 10a,b)**.

**Fig. 3.**
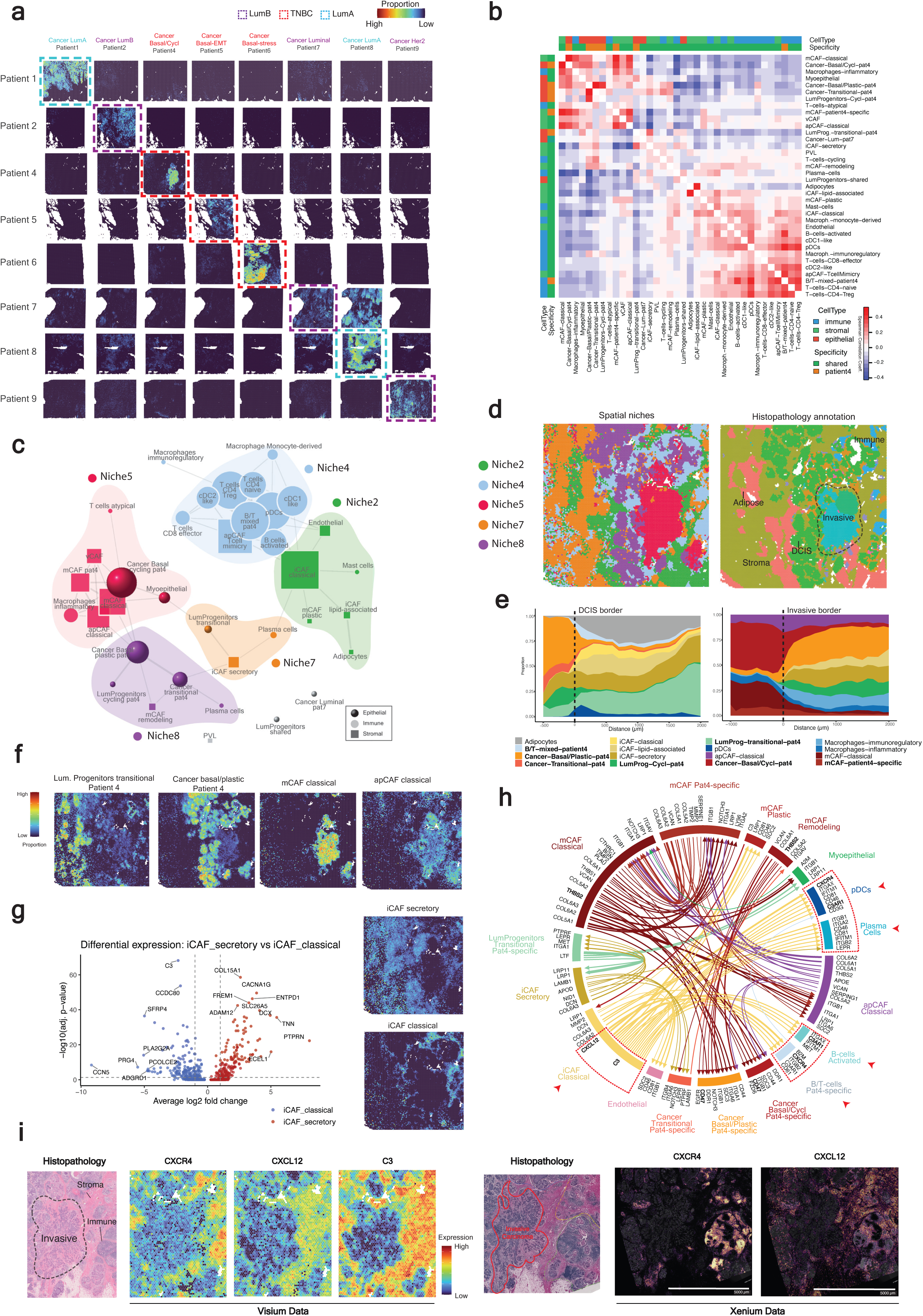
Mapping of high granularity cell states and patient-specific niche analysis. **a,** Spatial deconvolution mapping of SIMPlex cell states with one representative cell state for each section depicted. Dashed outlines denote BC molecular subtype. **b-i,** Exploration of Patient 4 signatures (**b**) Spatial co-localization analysis of patient-specific cell states. Heatmap shows Spearman Correlation Coefficient (SCC) of per-spot proportions. (**c)** Network representation cell state niches based on correlations obtained in (b). Edges represent spatial interactions established using a threshold of SCC > 0.15. Node size (circle/square/sphere) is proportional to the number of spatial associations. Niches were defined by louvain clustering of the network. (**d**) Spatial visualization cell state niches derived in (c) and histopathology. For each spot, the top-scoring niche label is displayed. (**e**) Cell state composition relative to DCIS and invasive borders. Stacked area plots show the top 10 cell states (patient-specific in bold) with the largest mean difference in proportion across the border, as a function of radial distance (µm). (**f**) Spatial mapping of key cell states demonstrating stromal and epithelial heterogeneity. (**g**) Volcano plot and spatial mapping of two iCAF populations. (**h**) Spatially informed ligand-receptor analysis. Circos plot visualizes the ligand-receptor pairs predicted from SIMPlex cell states that are spatially associated (SCC > 0.15) in (b) and significantly expressed (FDR < 0.05). Arrows connect sender cell state and ligand to receiver cell state and receptor, with arrow weight depicting interaction score. (**i**) Spatial validation of classical iCAF phenotype. Spatial maps display expression of genes selected based on ligand-receptor analysis (Xenium probes for C3 were not part of the panel). Histopathology on both confirms homologous tissue structures.

Niche-specific cell-cell communication was explored using spatially informed ligand-receptor analysis on matched nuclei data, filtering sender and receiver cell states by spatial proximity, to provide a contextual view of local microenvironment interactions (**Fig. 3h; Extended Data Fig. 10c,d**). iCAFs positioned in the tumor-immune interface featured *C3* and *CXCL12* secretion towards B cells and pDCs. The *CXCL12-CXCR4* axis has been widely linked to poor prognosis and immune suppression^27,28^. Despite associations in other cancers^29,30^, the role of *C3* in breast cancer is still underexplored. Same-section spatial transcriptomics and consecutive in situ profiling corroborate this peri-tumor arrangement, with both platforms showing *CXCL12* enrichment in the stroma neighboring *CXCR4+* B cells (**Fig. 3i; Extended Data Fig. 10e,f**). In contrast, non-malignant regions revealed reciprocal interactions between secretory iCAFs and patient 4-specific luminal progenitors^31–33^, while mCAFs in invasive/DCIS showcase interactions, such as *THBS2-CD47,* that have been associated with poor prognosis^34^.

Together, these results establish SIMPlex as a valuable framework for jointly profiling spatial and nuclear transcriptomes from a single tissue section, enabling molecularly resolved characterization of archival tissue. Although demonstrated in a modest cohort and tested on the Visium platform, SIMPlex reliably resolved both shared and patient-specific programs, revealing stromal and epithelial complexity; the approach could likely be extended to other spatial platforms as well. As a proof of concept, ligand-receptor analysis in a single section illustrated how matched data can reveal organised stromal-immune interactions, including potentially novel interactions at invasive borders. These capabilities create opportunities for spatially anchored biomarker discovery and precision-oncology research in clinical applications.

## Methods

### Samples and ethical permits

FF mouse brain tissue samples were purchased from Adlego Biomedical AB/Scantox, Sweden/Denmark. The FF breast cancer sample was purchased from BioIVT, UK. The FFPE blocks were sourced under “The Personalised Medicine for Breast Cancer Biobank” protocol (x19-0496 & 2019/ETH13373) approved by Sydney Local Health District (Royal Prince Alfred Hospital zone Committee, Sydney Australia). All patients provided informed consent for collecting and using their tissue in breast cancer-related research. The prostate cancer samples were obtained through open radical prostatectomies at Churchill Hospital in Oxford. All patients were provided with full verbal and written information about the study before their participation.

### Tissue preparation

FF mouse brain and FF breast cancer tissues were embedded in O.C.T. and stored in −80 °C until use. FFPE breast cancer sections were received at room temperature on the SuperFrost microscope slides.

### Mouse brain sample preparation

FF mouse brain sample was sectioned at 12 µm thickness. Two consecutive sections were placed within a pre-marked 11×11 mm area on each SuperFrost slide. Slides with sections were stored at −80°C prior to processing with the SIMPlex protocol.

### Breast cancer sample preparation

FF breast cancer samples were sectioned at 18 µm thickness. Each single section was placed into a pre-marked 11×11mm area on a SuperFrost slide and stored at −80°C prior to processing with the SIMPlex protocol. FFPE breast cancer samples were sectioned at 5 µm thickness, placed on Superfrost slides, stained according to clinical pathology workflows, hard coverslipped, imaged and shipped to our laboratory at room temperature. Slides were decoverslipped following an optimized version of the 10x protocol Visium CytAssist Spatial Gene Expression for FFPE - Tissue Preparation Guide (Document CG000518).

### Prostate cancer sample preparation

FFPE prostate cancer samples were prepared the same way as FFPE breast cancer samples.

### SIMPlex protocol

Part I: Visium spatial gene expression preparation (SIMPlex-Visium).

Fresh-frozen breast cancer and mouse brain tissue sections on SuperFrost slides were first incubated for 1 minute at 37°C on a thermal cycler and then fixed with 4% formaldehyde for 10 minutes at room temperature. Following the Hematoxylin and Eosin staining and imaging (10x Genomics, Document # CG000520). Next, slides were processed following the Visium CytAssist Spatial Gene Expression protocol (10x Genomics, Document # CG000495) for whole-transcriptome capture on Visium CytAssist slides. After RNA capture on the CytAssist instrument, the tissue slides were used for nuclei-isolation while the transcripts capture probes on the Visium CytAssist slides continued to extension, elution, library preparation and sequencing steps. The final Visium libraries were sequenced on the Nextseq2000 (Illumina) platform. The lengths of read 1 and read 2 were 28 and 50 base pairs, respectively.

Eight FFPE breast cancer tissue sections were stained and hard coverslipped according to pathology workflows, Coverslip removal (10x Genomics, Document #CG000518) and Decrosslinking protocol (10x Genomics, Document #CG000520) and processed on the Visium CytAssist. Two FFPE breast cancer tissue sections were prepared according to the protocols for Visium HD analysis from 10x Genomics, optimized version of Document CG000518 for Coverslip removal, Document # CG000684 for Decrosslinking step and Document # CG000685 User Guide. The final Visium HD libraries were sequenced on the Nextseq2000 (Illumina) platform. The lengths of read 1 and read 2 were 43 and 50 base pairs, respectively. All 10 FFPE breast cancer tissue sections were stored at −80 °C for the Part II of the SIMPlex protocol.

Part II: Nuclei Isolation and single nuclei library preparation (SIMPlex-snRNA).

The post-Cytassist tissue sections on SuperFrost slides were first re-fixed in 4% formaldehyde for 15 minutes at room temperature and washed with 1xPBS. Next, the sections were scraped with a cell scraper using 50 µl of nuclease-free water into a PCR tube pre-coated with 1xPBS + 2% BSA solution. Each sample was mixed with the probe hybridization mix, prepared following the 10x Genomics CG000477 protocol or CG000527 for Chromium Fixed RNA profiling, and incubated at 42°C overnight. The following day, 175 µl of Post-hybridization wash buffer was added to each sample, and samples were transferred to a 1.5 ml 1xPBS + 2% BSA pre-coated tube. The remaining Post-hybridization solution volume was added to reach a total volume of 900 µl. Subsequently, the samples were incubated for 10 minutes at 42°C and centrifuged at 850 rcf for 10 minutes at room temperature. This step was repeated, resulting in a total of two washes. Finally, the pellet was resuspended in 500 µl of Resuspension buffer. Next, an in-house nuclei isolation protocol was used to extract nuclei, during these steps all tubes were coated with 1xPBS + 2% BSA solution. Samples were placed on ice, and to each, 300 µL of lysis buffer (10 mM Tris-HCl, 10 mM NaCl, 3 mM MgCl2, 0.1% Igepal, 1 mM DTT, and 1 U/µL RNase Inhibitor) was added. Subsequently, the tissue in each sample was homogenized using a pestle. After homogenization, 500 µl of lysis buffer was added to the samples followed by incubation on ice. Of note, for mouse brain tissue, we did not perform any lysis step, while for breast and prostate cancer samples they were incubated for 12 and 15 minutes on ice, respectively. The samples were gently mixed several times with 1ml pipette during the incubation to facilitate lysis. The samples were first filtered through a 70 µm strainer to remove tissue debris, and the strainer was rinsed with 200 µl of lysis buffer to reduce sample loss. This process was then repeated with a 20 µm strainer, including the rinse step, to isolate nuclei further and reduce debris before the sorting. The samples were centrifuged at 500 rcf for 10 minutes at 4°C. Following centrifugation, each nuclei pellet was resuspended in 200 µL of 1x PBS + 2% BSA. After staining with DAPI, the samples were sorted using FANS (fluorescence activated nuclei sorting). A snRNA-seq sorting gate strategy was implemented to remove debris and doublets, recovering a singlet nuclei fraction, including 2n, 4n and fragmented 2n populations excluding the smallest nuclei fragments (Extended Data Fig. 3 b). The sorted nuclei were again centrifuged under the same conditions (500 rcf for 10 minutes at 4°C), resuspended in Post-Hybridization Resuspension buffer (10x Genomics) to achieve the required concentration and counted with a Countess II automated cell counter. The samples were loaded into Chromium X without further selection of specific nuclei populations aiming to profile 10,000 nuclei. Subsequent steps were conducted according to the 10X Genomics Chromium CG000527 user guide. The final libraries were sequenced on the Nextseq2000 (Illumina) platform, with Read 1 and Read 2 lengths of 28 and 90 base pairs, respectively.

### Step-by-step SIMPlex protocol

1. The SuperFrost slides, containing tissue sections were incubated for 1 minute at 37°C.
2. Sections were fixed with 4% formaldehyde (Thermofisher, Catalog number 28906) for 10 minutes at room temperature.
3. Slides were washed in 1xPBS (Medicago, Article number 09-9400). Then stained with Hematoxylin (Dako, Part number S330930-2) and Eosin (Sigma-Aldrich, Product number HT110216), followed by imaging.

I. *Generation of spatial libraries*.

*Next steps are performed according to the Visium Cytassist Spatial Gene Expression protocol (10x Genomics):*

4. The SuperFrost slides were placed into Visium cassettes and tissue sections were incubated with Pre-Hybridization Mix for 10 minutes at room temperature.
5. Pre-Hybridization Mix was replaced with Probe Hybridization Mix (for mouse and human) and incubated at 50°C overnight.
6. After performing the post-hybridization wash, the procedure was followed by probe ligation and a subsequent post-ligation wash.
7. Probe Release and hybridization to the Visium slide was performed using the CytAssist machine.
8. Following the Cytassist step, the Visium slide was placed into a cassette and incubated with Probe Extension Mix.
9. Probes were eluted and each sample was collected into a tube for pre-amplification step.
10. Samples were cleaned using SPRIselect beads (Beckman Coulter, Product number B23318) and indexed via PCR reaction.
11. Samples underwent a second cleanup using SPRIselect beads and the measurement of the concentration and length of the final libraries were performed prior to the sequencing using Qubit (Thermo Fisher, Product number Q33230) and Bioanalyzer 2100 (Agilent High Sensitivity DNA Kit, Product number 5067-4626) respectively.

II. *Generation of single nuclei libraries*.

12. Tissue sections on SuperFrost slides that underwent CytAssist run were re-fixed using 4% formaldehyde for 15 minutes at room temperature.

Note: FFPE-BC samples were processed approximately six months after the ST experiments. The slides were stored in a −80°C freezer until the time of nuclei scraping and isolation.

13. Slides were washed with 1xPBS and 50 µl of nuclease free water was added on top of each section, which was subsequently scraped into a PCR tube pre-coated with 1xPBS + 2% BSA.
14. Each sample was resuspended in 80 µl of the Hybridization Mix.
15. Then, 20 µl of RNA probes was added to the sample, pipette-mixed and incubated at 42°C overnight.
16. Samples were washed and centrifuged at 850 rcf for 10 minutes at room temperature twice.
17. The pellet was resuspended in 200 µl of chilled Resuspension buffer.

*Next steps are performed according to the in-house nuclei extraction protocol:*

18. 300 µl of Lysis buffer (10mM Tris-HCl, 10mM NaCl, 3mM MgCl2, 0.1% Igepal, 1mM DTT, 1U/µl RNase Inhibitor) was added to the pellet and homogenized with a pestle (SP Bel-Art, Catalog number 199230001).
19. 500 µl of lysis buffer was added to the sample and incubated on ice for 12 min (only for breast cancer and 15 min for prostate tissues and no lysis for mouse brain. Mouse brain sample was dissociated in lysis buffer by pipetting without additional incubation time). Sample was mixed several times during the incubation using 1 ml pipette.
20. Samples were passed through a 70 µm strainer (pluriSelect, SKU 43-50070-51) into a new coated tube (1xPBS + 2% BSA). Strainer was washed with 200 µl of lysis buffer to reduce the loss of nuclei.
21. Samples were passed through a 20 µm strainer (pluriSelect, SKU 43-50020-03) and washed with 200 µl of lysis buffer.
22. Samples were centrifuged at 500 rcf at 4°C for 10 min and pellet containing nuclei was resuspended in 200 ul of 1xPBS + 2% BSA, stained with DAPI (Thermo Scientific, Product number 62248) and sorted by flow cytometry, utilizing a BD FUSION Aria III sorter.
23. After sorting, samples were centrifuged at 500 rcf for 10 minutes at 4°C.
24. Pellets containing nuclei were resuspended with Post-Hybridization Resuspension buffer and counted using Countess II automated cell counter, and loaded into the Chromium X system. *Next steps are performed according to the Chromium Fixed RNA Profiling protocol (10x Genomics):*
25. Recovered GEMs were incubated in a thermocycler according to the protocol, followed by GEM recovery and Pre-Amplification PCR.
26. Samples were cleaned using SPRIselect beads, indexed via PCR reaction, followed by additional cleaning with SPRIselect beads.
27. Concentration and length of the final libraries were measured prior to the sequencing.

### Experimental considerations

Tissue sections met the 10x Genomics recommendations for RNA quality, with DV200 values >30% and RIN values >7. The tissue area analyzed ranged from 1 × 1 cm to 2 × 2 cm considered large tissue sections and delivering the nuclei recovery ranges documented in this work. Nuclei isolation relied on inexpensive standard laboratory reagents and access to flow cytometry. The main costs were associated with the Visium and Chromium Flex kits, while the risk of failure was low if maximizing tissue input and loading of debris-free preparations of single-sorted nuclei to the Chromium instrument. The recovery rates, DV200, estimated tissue area, and patient metadata for each sample are presented in Supplementary Table 1.

### Single-nuclei RNA sequencing experiment control

FFPE breast cancer tissue sections 5 µm thick (one from patient 4, three for patient 6) placed on slides were rehydrated and deparaffinized and scrapped according to Appendix A in 10x Genomics protocol CG000632 Rev D. Nuclei were isolated as in SIMPlex protocol *Part II: Nuclei Isolation and single nuclei library preparation.* The Visium experiment or *Part I: Visium spatial gene expression preparation* was completely omitted for this experiment. Isolated nuclei were hybridized with Flex probes as in 10x Genomics protocol CG000527 Rev D. Next day, nuclei were sorted according to our customized snRNA-seq sorting gate strategy and counted with a Countess II automated cell counter. The samples were loaded into Chromium X aiming to profile 10,000 nuclei. The final libraries were sequenced on the Nextseq2000 (Illumina) platform, with Read 1 and Read 2 lengths of 28 and 90 base pairs, respectively.

### Fluorescence activated nuclei sorting (FANS) to profile nuclei integrity

We assessed nuclei obtained from four human breast cancer FFPE sections samples of varying thicknesses in order to profile the changes in nuclei integrity. The nuclei were isolated from sections of 5, 10, 20 and 30 µm after de-paraffinization and following the nuclei isolation protocol as described in the manuscript with a modification on protease-based lysis for the thicker sections. The nuclei homogenates from each sample were spiked with pre-sorted intact nuclei stained with 7-AAD from a control sample that is intended for internal integrity and size reference. The control sample was prepared from a 30 μm-thick tissue section, the same human breast cancer FFPE tissue block as the four samples, homogenised as described in the nuclei isolation protocol in the manuscript with a modification on protease-based lysis and then stained with 7-AAD. The 7-AAD DNA stain was chosen in order to track the control intact 2n nuclei after spiked in the samples. The 7-AAD stained control sample was analysed with FANS and all the 2n single nuclei were sorted. A total of approximately 2.8·10^5 nuclei were collected. From the collected control 2n nuclei, 50·10^3 were spiked-in to each of the four sample homogenates and then stained with DAPI in order to have even DAPI staining between control and sample nuclei for robust quantitative analysis. The spiked samples were analysed with FANS. The nuclei integrity assessment was based on nuclei DNA content as measured by DAPI and 7-AAD, initially by comparing the nuclei isolates DAPI signal distributions and then by validating the target sub-populations by confocal microscopy. Images of sorted nuclei from the control sample and from sample sub-populations, intact and fragmented, were acquired. When we compared the distributions of 5, 10, 20 and 30 μm sections samples, we observed a gradual increase in the frequency of the truncated nuclei and a decrease in their size with decreasing section thickness as expected. Despite that, most of the nuclei fragments appear crisp with no signs of degradation or deformation and moreover intact 2n and 4n nuclei exist in the lowest thickness tissue section sample.

### Data pre-processing

Chromium single-nuclei FASTQ files were processed with Cell Ranger (version 7.1.0, 10x Genomics) using the 10x Genomics pre-built 2020-A reference (GRCh38, version 32, ensembl 98) and Chromium Mouse Transcriptome Probe Set v1.0.1 or Chromium Human Transcriptome Probe Set v1.0.1 (10x Genomics). The corresponding Visium FFPE CytAssist FASTQ files were processed with Space Ranger (version 2.0.1, 10x Genomics) using the same pre-built reference and the Visium Mouse Transcriptome Probe Set v1.0 or Visium Human Transcriptome Probe Set v2.0 (10x Genomics). Background noise such as ambient RNA was removed from the count matrix using the remove-background function of *CellBender*^35^) with default settings.

### Data filtering and processing

Single-nuclei data processing and visualization were carried out using the R^36^ (v 4.3.3) package *Seurat*^37^ (v 5.2.1) and *semla*^38^ (v 1.3.1) was used for spatial transcriptomics data processing, visualization, and cell type deconvolution.

### Data analysis for mouse brain samples

Two capture areas, each containing two mouse brain tissue sections, were used for the analysis, along with publicly available single-cell mouse forebrain FFPE data (10x Genomics). All the generated mouse brain data were analyzed jointly. Violin plots of unique genes and UMI counts for both single-nuclei and spatial data were visualized using the VlnPlot() function (*Seurat*). Gene-gene scatter plots comparing log-transformed UMI counts were created by log-transforming aggregated expression values for each gene obtained by extracting raw expression matrices for each sample. Pearson R scores and p-values were calculated using the stat_cor() function from the *ggpubr*^39^ (v 0.6.2) R package. Gene-gene scatter plots comparing detection rates were created by extracting raw expression matrices for each sample followed by estimation of detection rate for each gene as the proportion of spots with detected UMI counts. Generated single-nuclei data was compared to a publicly available mouse forebrain dataset (10x Genomics), while the spatial data was compared between two technical replicates. Genes with fewer than 3 UMIs across all nuclei were filtered out, while the criteria for the spatial spots were at least 5 UMIs per spot and more than 300 UMIs across spots. Nuclei and spots with fewer than 400 genes were also filtered out in both datasets. After filtering, the data was normalized and subjected to a basic analysis workflow using functions from the *Seurat* R package.

#### Nuclei data

Normalization was performed using SCTransform() with the variance stabilizing transformation (vst) set to v2 followed by dimensionality reduction by RunPCA(). Doublets identification and removal were carried out using *DoubletFinder*^40^ (v 2.0.3) by first finding the optimal pK value without ground-truth, computed by paramSweep and summarizeSweep using the first 30 principal components, followed by doublet identification through doubletFinder_v3 with pN set to 0.25 and expected doublets set to 10%. Following doublet removal, normalization was performed once again, followed by scaling and dimensionality reduction using SCTransform() with vst set to v2, ScaleData() and RunPCA(). A Uniform Manifold Approximation and Projection (UMAP) embedding was computed based on the first 30 principal components with default settings (RunUMAP). Annotation was carried out by label transfer from publicly available single-cell mouse brain data sets^15,16^ using the FindTransferAnchors() and TransferData() functions with default settings. Marker detection was conducted by calculating differential expression for each cluster against the background (remaining clusters) with a log fold change threshold of 0.25 and an adjusted p-value threshold of 0.01 using the FindAllMarkers() function. Additionally, only genes detected in 0.25% of either of the two populations were tested to speed up the process, with the top 5 marker genes visualized using the DotPlot() function.

Spatial data: Spatial transcriptomics data were analyzed using functions from the *semla* R package^38^. Normalization was performed using NormalizeData() followed by detection of the top 10,000 most variable genes using the vst method (FindVariableFeatures()). Before deconvolution, FindVariableFeatures() was repeated on the single-nuclei data, selecting for the top 10,000 most variable genes, to increase the number of genes before cell type mapping. The RunNNLS function was used to infer the quantity of cells from spatial transcriptomics data with nCells_per_group set to 8000, 4000, and 2000 for subclass, taxonomy4, and Allen cortex-derived single-nuclei annotations, respectively. The distribution of the inferred cell type quantities was visualized on top of the histology image using the MapFeatures() and MapMultipleFeatures() functions for single cell types or multiple cell types using the image_use parameter.

### Data analysis for breast/prostate cancer

#### Nuclei data processing

Data was processed per-sample with a similar workflow of doublet removal as described for the mouse samples. Nuclei with fewer than 200 unique genes and genes with fewer than 3 UMIs across all nuclei were removed. Prior to data merging, on individual nuclei samples before cell type assignment, data was normalised with Seurat’s SCTransform() (v2 flavor), Principal Component Analysis (PCA) with 2000 most variable features and default 50 dimensions using Seurat’s RunPCA(), and UMAP visualization with 30 dimensions via RunUMAP().

#### Cell type assignment

Nuclei were assigned cell type identities through an iterative process combining reference-based label transfer and marker gene inspection. Initial labels were obtained by Seurat’s (version here) FindTransferAnchors() and TransferData() using the breast cancer single-cell atlas from Wu *et al.*^3^ as reference. Per-sample clusters were then relabelled based on the transferred labels and marker gene inspection. Confirmation, merging, or subdivision of transferred labels and clusters were guided by visualization of lineage-defining markers across adipocyte *(PLIN1, ADIPOQ, LEP, PPARG, FABP4), B cell (MS4A1, CD19, CD79A, CD79B, CD22),* T cell *(CD3D, CD3E, CD3G, CD4, CD8A, CD8B, PDCD1),* plasmablast *(JCHAIN),* endothelial *(PECAM1, CDH5, VWF), PVL (PDGFRB, MCAM, CD146, ACTA2), myeloid (CD68, CD1C), CAF (VIM, COL1A1, PDGFRB, ACTA2),* and epithelial *(EPCAM, FOXA1)* populations.

#### Data integration

After annotation of each individual sample, nuclei data was compiled into a single object and analyzed using Seurat’s standard approach and integrated using Harmony^41^ (v 1.2.3). Merged data was processed using Seurat’s standard log-normalization workflow (NormalizeData(), FindVariableFeatures() (2000 features), ScaleData()). PCA was run with default 50 components. To retain biological diversity while correcting batch effects we ran Harmony (specifying “sample” as covariate) using a theta of 0 to avoid over-correction. A UMAP embedding was computed both before and after integration (using 30 first dimensions) to evaluate batch effects, cell type structure and biological signal preservation. Identification of Cancer *versus* tumor epithelial populations was done via label transfer (as described previously) from Wu *et al.*^3^.

#### Cell type mapping

We used NNLS^42^ implementation in the *semla*^38^ package to perform cell type mapping to standard and HD Spatial Transcriptomics data. For each Visium sample, we used Technical/Pooled SIMPlex controls, BC reference atlas, and SIMPlex data to map cell types using the RunNNLS() function in Semla. Top 2000 variable features were selected via FindVariableFeatures() for the source single nuclei/cell data and all features in Visium data were used. Concordance of mapping between different references and to marker genes was computed using Spearman correlation and visualized using heatmaps using the heatmap3^43^ (v 1.1.9) R package. Lineage-specific module scores were calculated using UCell^44^ (v 2.6.2) with a list of marker genes for B cells (*CD79B*, *MS4A1, BANK1*), T cells (*CD3D, CD3E, CD3G, CD2, TRAC*), plasmablasts (*XBP1, PRDM1, JCHAIN, IGHG1*), myeloid cells (*ITGAM, MRC1, CD68, FUT4, FCGR3A*), epithelial cells (*EPCAM, KRT8, KRT5, KRT14, CDH1, MUC1, CLDN3, CLDN4, CLDN7)*, fibroblasts (*COL1A1, COL1A2, DCN, FAP, MMP2, MMP11, THY1*), endothelial cells (*PECAM1, VWF, CDH5, ESAM, TIE1, TEK, FLT1, KDR, CLDN5, PLVAP, PROX1, LYVE1, PDPN*), and perivascular-like cells (*MCAM, ACTA2, PDGFRB*). Pairwise comparisons of per-cell type correlations between references were evaluated using Wilcoxon signed-rank tests and multiple testing was corrected with the Benjamini-Hochberg method.

#### CTA annotation

CTA annotation^18^ was performed using QuPath^45^ (v 5.0.1) on the H&E images in TIFF format. After stain vector correction, cell segmentation was applied through the Cell detection function, followed by calculating smoothed features (Add smoothed features function) and training object classifier. The annotated images were aligned back to the Visium spots using the CTA_align function. We compared spot-level CTA annotation classifications to deconvolution using the different references by computing Spearman correlation and visualized as heatmaps (heatmap3 (v 1.1.9)) and boxplots.

#### Spatial gene imputation

Visium HD count matrices (8 µm bins; Patient 5) were imputed with SpaGE^46^ using either section-matched SIMPlex data, pooled SIMPlex data, section-unmatched SIMPlex data (all other sections), BC Atlas, and patient-matched BC Atlas (CID44971 for Patient 5). Analysis was performed on a central tissue ROI (as in Fig. 2i). Spatial and sn/scRNA matrices were log-normalized and restricted to shared genes in both datasets, followed by Z-scoring per gene and PCA. SpaGE was run with 30 principal vectors with cosine similarity above 0.3 and 50 nearest neighbours. Imputation was performed separately per reference. A predefined set of markers were imputed (as per UCell module scoring) and visualized spatially.

#### Annotation of fine-grained cell states in SIMPlex data

Subclustering and annotation of Epithelial, CAF, and Immune populations was done with a combination of manual, comparative and label transfer approaches. Cross-patient Epithelial, CAF, and immune populations from the integrated dataset were subset from the original object and filtered in order to proceed with high-confidence populations only. CAFs were selected with an expression threshold of 0.5 for positive (*COL1A1* and *COL1A2*) and negative (*EPCAM, KRT8, CD3D, CD3E, MS4A1, TRAC, CD163*) markers. Immune populations were initially filtered by label transfer with Seurat’s FindTransferAnchors() and TransferData() from Wu *et al*.^3^ reference and elimination of possible epithelial/fibroblast contamination of nuclei with expression *EPCAM* and *COL1A1* larger than 1. For all lineages, after subsetting from the original object, data was normalized using NormalizeData() (default parameters), highly variable genes identified with FindVariableFeatures, and data scaled with ScaleData(). Pre-integration PCA was run with RunPCA(), integration using RunHarmony() (theta = 0) for the Patient ID covariate, and RunUMAP() using the first 30 dimensions using “harmony” dimensionality reduction. Post-integration clustering was performed using FindNeighbors() with the first 30 dimensions, and clustering with FindClusters() at the different resolutions of 0.1, 0.25, 0.3, 0.4, 0.5, 0.8, 1.0, 1.25. The following resolutions were used for each of the lineages: CAFs (resolution = 0.8); Epithelial (resolution = 1); Immune (resolution = 0.8). We compared our lineage subclusters to relevant literature and generated correlation heatmaps to inform our annotation decisions. Wu *et al.*^3^ “minor” populations were used for Immune and Epithelial populations and Cords *et al.*^22^ for CAF populations. Top 3000 features were picked in each of the lineages and compared to the per-gene population averages in the reference by using Seurat’s AverageExpression and computing per-population Pearson correlation coefficients. This was displayed in a heatmap (heatmap3 (v 1.1.9)) in order to find clusters in our data that relate to reference populations. Label transfer (as described above) was also used to compare our clusters to the corresponding atlases. To inform our final classification, cluster-specific markers were obtained by Seurat’s FindAllMarkers() and exported the top 100 markers per cluster for further investigation. Additionally we used Gene Set Enrichment Analysis to investigate pathways represented in our clusters by using the R package singleseqgset^47^ (v 0.1.2.9000) with hallmark gene sets from MSigDB (H category; MSigDB v7.5.1 release) accessed via the msigdbr^48^ (v 7.5.1) package in R.

#### Spatial co-analysis of SIMPlex cell states

Visium cell type deconvolution with SIMPlex data was performed as described above using RunNNLS() with 2000 top variable features in Visium data, but now selecting top 2000 variable features in the reference as well (for specificity). Two modes of deconvolution using SIMPlex were used. For evaluating the mapping of patient-specific single-nuclei signatures to its cognate Visium section, the pooled dataset including all patients was used, where we can assess all populations across all Visium data. In contrast, to explore patient-specific biology and avoid dilution of patient-specific signatures in shared profiles (such as CAF and Immune populations) we use each patient’s single nuclei data to deconvolve each corresponding Visium section. Global, subtype-, and patient-specific correlations of per-spot deconvolved cell states were computed generating Spearman correlation coefficients. When visualizing global or subtype-specific correlations, shared biological terms were collapsed into the same category (e.g. patient4 and 5-specific basal cancer into “basal cancer”) to avoid signal dilution. Per-patient correlation matrices were used to compute colocalization networks using the igraph^49^ (v 2.1.4) R package following the implementation applied in Lázár *et al.*^25^. Cell state pairs with Spearman correlation scores below the arbitrary threshold of 0.15 were excluded in order to preserve spatially relevant interactions, graph_from_adjacency_matrix() function used to generate the graph (weighted = TRUE, mode = “undirected”), degree() function to calculate the number of edges to each vertex, and Fruchterman-Reingold algorithm used for layout with the layout_with_fr() function. Niches were identified by applying a Louvain clustering algorithm using cluster_louvain() with a resolution of 1.7. Per-spot niche scores were calculated by summing constituent cell state proportions of each niche followed by Z-score normalization and stored in each object as “niche_score”. Niches were visualized spatially by labelling each spot by its highest scoring niche. As a comparison, deconvolution with Wu *et al.*^3^ was performed using the fine-grain population levels “celltype_subset” with Semla’s RunNNLS() (with the same parameters as SIMPlex data).

#### Cluster stability analysis

Fine-grained subpopulations were tested within CAF, epithelial, and immune compartments using *scclusteval*^50^. We downsampled to 80% of the original nuclei in 20 iterations. Nuclei were subsampled, reclustered (using same parameters as in the original analysis), and each original subpopulation was matched to its best subsample cluster. Stability was quantified as the median Jaccard index across iterations (stable ≥ 0.6; highly stable ≥ 0.85).

#### Spatial niche threshold sensitivity

Per-sample Visium sections were analysed using deconvolved subpopulation proportions per spot. Pairwise Spearman correlations between subpopulations were computed as described before and converted into undirected weighted networks by retaining edges with ρ ≥ T, for thresholds T from 0 to 0.4 (in 0.005 increments). At each threshold, communities were defined by Louvain clustering (igraph^49^). Stability of the resulting niche partition was assessed by comparing each test partition to the reference partition obtained at r = 0.15 using the adjusted Rand index (mclust::adjustedRandIndex). Network size metrics (edge number, cluster number, cluster size) were also recorded across thresholds.

#### Spatial radial analysis

In Patient 4 standard Visium data, areas annotated as DCIS and Invasive were used to compute radial distances by using Semla’s RadialDistance() function (convert_to_microns = TRUE), assigning per-spot distances from the border of each selected area. We then visualized variation of cell states proportions across each respective border in stacked area plots in 100 μm distance bins. For each bin, we displayed the top 10 cell states showing the largest spot-averaged difference in proportion across the border in order to facilitate interpretation. Plasma cells were excluded from this plot due to possible over-representation issues in Visium data.

#### Differential expression analysis

Pairwise differential expression analysis between groups was performed using Seurat’s FindMarkers() (Wilcoxon rank-sum test; log2FC threshold = 0.25; minimum detection fraction = 0.1). For each comparison, gene-level statistics (log2 fold-change and FDR-adjusted p-values) were used to generate Volcano plots using the ggplot2^39^ (v 3.5.2) R package.

#### Spatially informed ligand-receptor analysis

Ligand-receptor interactions were inferred from single-nuclei data using the celltalker^51^ (v 0.0.7.9000) R package using ligand-receptor pairs from Ramilowski *et al.*^52^. Interactions were computed using the celltalk() function (number_cells_required=30, min_expression=200, max_expression=20000, scramble_times=10). Spatial constraints were introduced by incorporating previously obtained pairwise cell state Spearman correlation matrices. celltalker interaction statistics for sender and receiver cell states if corresponding Spearman correlation coefficients over 0.15 and FDR-adjusted p-values over 0.05. CCPlotR^53^ (v 1.0.0) R package was used to visualize interactions with the following functions: cc_heatmap(), cc_dotplot(), and cc_circos().

#### Xenium in situ profiling and annotation

Consecutive FFPE tissue sections adjacent to those used for Visium or Visium HD profiling were processed for independent in situ validation. Five-micrometer sections were analyzed using the 10x Genomics Xenium In Situ platform following the manufacturer’s instructions. The breast-cancer–targeted Xenium probe set was supplemented with 100 custom probes covering epithelial, stromal, and immune lineage markers (include *KRT7/8/5/6/14, ERBB2, PGR, MKI67, CD3E/G,* and *CD79A/B*). Raw Xenium outputs were decoded and segmented using the Xenium Analyzer pipeline to obtain cell-segmentation masks and per-cell transcript count matrices. Transcripts were assigned to cell boundaries, and cells failing basic quality filters (*e.g.*, low total counts or excessive background) were excluded. Per-cell counts were log-normalized prior to downstream analysis and visualization.

For cell type annotation, Xenium per-cell expression profiles were mapped to a breast-cancer single-cell reference^3^ using a two-stage label-transfer pipeline implemented in Python (Scanpy v 1.9). First, shared variable genes were selected between the reference and query datasets, and the query data were normalized and log-transformed. The reference atlas was embedded via PCA, neighborhood graph construction, and UMAP. Using scanpy.tl.ingest, major lineage identities were transferred from the reference to the Xenium dataset in a supervised manner. Subsequently, within each major lineage subset, local classifiers were retrained using the corresponding subset of the reference (*e.g*., epithelial, stromal, immune), and minor subtype labels were transferred iteratively using the same ingest framework. The resulting annotated dataset combined all lineage-specific predictions into a unified AnnData object. Predicted labels were further inspected and refined by overlaying canonical marker expression on H&E/Xenium composite maps, and low-confidence assignments were manually curated.

## Data availability

Processed data are deposited in Zenodo, DOI: 10.5281/zenodo.21497736, including CellRanger/CellBender and SpaceRanger outputs, processed Xenium data, annotated Seurat objects, histopathology tables, and figure files for breast cancer, prostate cancer, and mouse brain cohorts.

Raw Chromium snRNA-seq and Visium FASTQs, and source H&E/CytAssist microscopy where applicable, are deposited separately in [REPOSITORY TBD] under accession [ACCESSION TBD]. Public reference datasets used for benchmarking and annotation are documented in the GitHub repository (docs/data_availability.md).

## Code availability

Analysis code is available at https://github.com/spatial-research/SIMPlex_analysis, including Jupyter notebooks, environment files (environment/setup.sh, environment/renv.lock, environment/environment.yml). Processed data from the Zenodo archive can be unpacked into the repository root to regenerate the figures. The specific version used for the analyses presented in this manuscript is archived in Zenodo (DOI: [DOI TBD]).

## Acknowledgements

This project has received funding from the European Research Council (ERC) under the European Union’s Horizon 2020 research and innovation programme (grant agreement no. 101021019 J.L.). The study was also supported by The Swedish Cancer Society (grant agreement 71170; 2024-01936 J.L.), a grant from the Breast Cancer Research Foundation USA (BCRF-24-209), Swedish Foundation for Strategic Research (grant agreement SB16-0014, J.L.), a Clinical Extension Award from the US Government CDMRP breast cancer program (RG242379), the Swedish Research Council (Dnr: 2022-03984; 2024-01936 J.L.) and Science for Life Laboratory (J.L.). A.S. is supported by an Investigator grant from the National Health and Medical Research Council, Australia (APP2018440) and by the generosity of the Skipper-Jacobs Charitable Trust, The Petre family, John McMurtrie, AM and Deborah McMurtrie. We would like to thank National Genomics Infrastructure (NGI), Sweden, for providing infrastructure support. We thank Drs. Ludvig Larsson, Enikö Lázár, Hani Kim and Eva Gracia Villacampa for helpful assistance.

## Authors and Affiliations

**Department of Gene Technology, KTH Royal Institute of Technology, Science for Life Laboratory, Stockholm, Sweden**

Marcos Machado, Mengxiao He, Leire Alonso Galicia, Zaneta Andrusivova, Raphaël Mauron, Javier Escudero Morlanes, Marco Vicari, Emmanouela Perisynaki, Joakim Lundeberg, Reza Mirzazadeh

**National Institute of Oncology, Budapest, Hungary**

Máté Mihálffy

**Department of Cell and Molecular Biology, Karolinska Institute, Stockholm, Sweden**

Sarantis Giatrellis

**Cancer Ecosystems Program, Garvan Institute of Medical Research, Darlinghurst, NSW 2010, Australia**

Beata Kiedik, Kate Harvey, Taopeng Wang, Alex Swarbrick, John Reeves

Dame Roma Mitchell Cancer Research Laboratories, Adelaide Medical School, University of Adelaide, Adelaide, South Australia, Australia

Beata Kiedik

**School of Clinical Medicine, Faculty of Medicine and Health, UNSW Sydney, Australia**

Taopeng Wang, Alex Swarbrick

**iCAN Digital Precision Cancer Medicine Flagship, University of Helsinki and Helsinki University Hospital, Helsinki, Finland**

Andrew Erickson, Laura Savolainen

**Department of Biochemistry and Biophysics, Stockholm University, Science for Life Laboratory, Stockholm, Sweden**

Mats Nilsson, Mengping Long, Taobo Hu

**Department of Oncology-Pathology, Karolinska Institutet, Stockholm, Sweden**

Tianyi Li, Xinsong Chen, Johan Hartman

**Department of Clinical Pathology and Cancer Diagnostics, Karolinska University Hospital, Stockholm, Sweden**

Johan Hartman

**University of Oxford, Nuffield Department of Surgical Sciences, Oxford, United Kingdom**

Sandy Figiel

**Centre for Cancer Evolution, Barts Cancer Institute, Queen Mary University of London & Department of Urology, Guy’s Hospital, London, UK**

Alastair Lamb

## Author contribution

R.M. and J.L. initiated the project and together with M.T.M., M.H., L.A.G., Z.A. designed and planned the study and analysis. L.A.G., R.M., Z.A., M.H., E.P, M.W.M performed the experiments. M.T.M., performed BC and PCa spatial and nuclei analysis with initial help from M.H. M.H performed Mouse spatial and nuclei analysis. M.T.M and B.K. performed BC single-nuclei annotation and interpretation. S.O., M.M. and M.L. annotated histology samples. L.S., A.E. helped histology annotation of fresh frozen samples. T.L., X.Ch. and J.H. performed CTA annotations. T.H. and M.N. performed Xenium experiments. K.H., T.W., S.L., J.R., A.S. provided FFPE-BC specimens, biological insights and feedback. S.G. led the nuclei sorting part and their optimization. S.F. and A.L. provided FFPE-prostate sections. M.V., R.Ma and J.E.M. helped in the initial phase of the project. R.M., M.T.M, M.H., L.A.G. drafted the paper with input from all the authors. R.M. and J.L. provided project guidance and supervision.

Correspondence to Joakim Lundeberg.

