## Supplementary Table 1 for "Highly resolved tumor architecture via matched spatial and nucleus transcriptomics from a single tissue section"

| sample_id | patient_id | preservation_m | subtype | collection_year | tissue_area | spatial_assay | dv200_rin | pools |
|---|---|---|---|---|---|---|---|---|
| MB_2 | MB_2 | FF | Mouse Brain | NA | 0.5 x 0.5 cm | Visium | NA | 1 |
| patient10_55un | patient10 | FF | unknown | 2012 | 1 x 1 cm | Visium | 8.1 | 2 |
| patient10_ctrl1 | patient10 | FF | unknown | 2012 | 1 x 1 cm | NA | 8.1 | 1 |
| patient10_ctrl2 | patient10 | FF | unknown | 2012 | 1 x 1 cm | NA | 8.1 | 1 |
| patient1_55um | patient1 | FFPE | LuminalA | 2014 | 2 x 2 cm | Visium | NA | 1 |
| patient2_55um | patient2 | FFPE | LuminalB(ER) | 2016 | 2 x 2 cm | Visium | NA | 1 |
| patient4_55um | patient4 | FFPE | TNBC | 2015 | 2 x 2 cm | Visium | 52 | 2 |
| patient4_HD | patient4 | FFPE | TNBC | 2015 | 2 x 2 cm | Visium HD | 52 | 2 |
| patient4_ctrl | patient4 | FFPE | TNBC | 2015 | 2 x 2 cm | NA | 52 | 1 |
| patient5_55um | patient5 | FFPE | TNBC | 2018 | 1 x 1 cm | Visium | 44 | 2 |
| patient5_HD | patient5 | FFPE | TNBC | 2018 | 1 x 1 cm | Visium HD | 44 | 1 |
| patient6_55um | patient6 | FFPE | TNBC | 2020 | 2 x 2 cm | Visium | 49 | 2 |
| patient6_ctrl_3: | patient6 | FFPE | TNBC | 2020 | 2 x 2 cm | NA | 49 | 1 |
| patient7_55um | patient7 | FFPE | LuminalB(ER) | 2014 | 2 x 2 cm | Visium | NA | 1 |
| patient8_55um | patient8 | FFPE | LuminalA | 2021 | 2 x 2 cm | Visium | NA | 1 |
| patient9_55um | patient9 | FFPE | LuminalB(ER) | 2016 | 2 x 2 cm | Visium | 51 | 1 |
| pt10_HD | pt10 | FFPE | Prostate Cance | 2019 | 1 x 1 cm | Visium HD | NA | 1 |
| pt20_HD | pt20 | FFPE | Prostate Cance | 2019 | 1 x 1 cm | Visium | NA | 1 |
| median_global | | | | | | | | 1 |
| median_BC | | | | | | | | 1 |
| median_mouse | | | | | | | | 1 |
| median_PC | | | | | | | | 1 |
| legend | Clinical metada | Tissue preserv: | Molecular subty | Year of tumour | Approximate ar | Spatial transcri | RNA quality me | Number of snR |

| cellbender_nuc | final_nuclei | median_genes | median_counts |
|---|---|---|---|
| 14575 | 11129 | 2747 | 5047 |
| 69435 | 48464 | 1025 | 1596 |
| 6497 | 5917 | 950 | 1400 |
| 6571 | 5995 | 1646 | 2748 |
| 19679 | 10495 | 636 | 788 |
| 13018 | 8401 | 851 | 1212 |
| 4165 | 3049 | 600 | 710 |
| 10379 | 7751 | 789 | 1080 |
| 17483 | 9135 | 168 | 185 |
| 9860 | 7678 | 798 | 950 |
| 12362 | 6709 | 633 | 765 |
| 9105 | 7229 | 1039 | 1466 |
| 25869 | 17718 | 171 | 207 |
| 8378 | 5653 | 493 | 547 |
| 5834 | 4990 | 747 | 901 |
| 11161 | 9463 | 1154 | 1462 |
| 6475 | 5659 | 625 | 726 |
| 9905 | 7734 | 377 | 439 |
| 10142 | 7706 | 768 | 926 |
| 10379 | 7678 | 789 | 950 |
| 14575 | 11129 | 2747 | 5047 |
| 8190 | 6696 | 501 | 582 |
| Nuclei called by | Nuclei retained | Median genes ( | Median UMIs per nucleus (nCount_RNA) in the final object. |
