## Extended Data Fig. 10 for "Highly resolved tumor architecture via matched spatial and nucleus transcriptomics from a single tissue section"

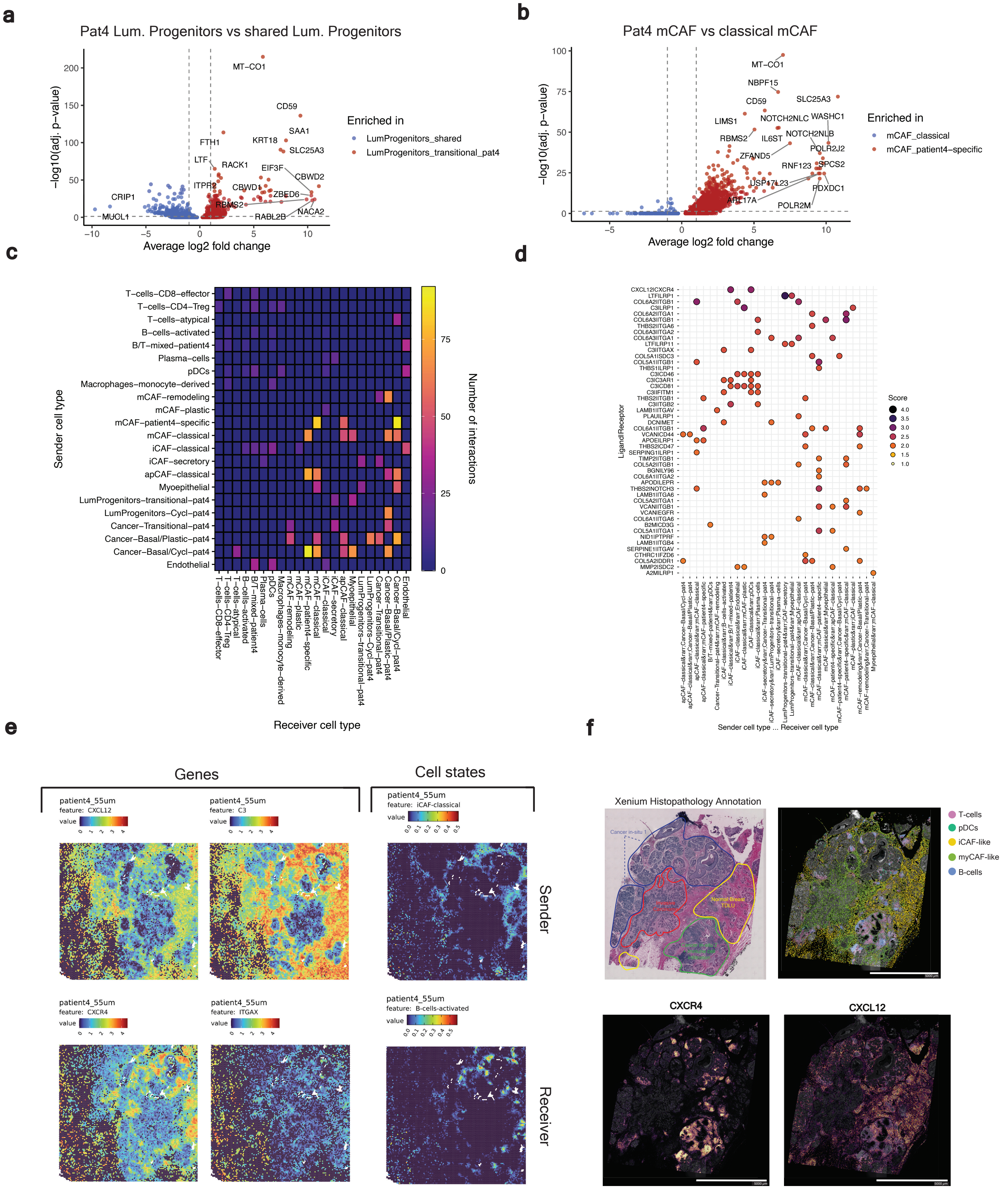

**Extended Data Fig.10 | Exploration of stromal signatures in patient 4 and cell-cell communication. a,b,** Volcano plot showing differential expression between key epithelial and stromal populations. (a) Compares Patient 4-specific Luminal Progenitors to the remaining patient-shared Luminal Progenitor population. (b) Compares Patient 4-specific mCAF population to classical mCAF population shared across patients. **c,d,** Spatially informed ligand-receptor analysis in Patient 4 with ligand-receptor pairs (see methods) predicted from SIMPlex cell states that are spatially associated (SCC > 0.15) and significantly expressed (FDR < 0.05). (c) Heatmap displaying number of positive interactions between cell states. (d) Dot plot displaying ligand-receptor score for specified sender and receiver cell states. **e,** Spatial visualization of Visium gene expression data for CXCL12, C3, and CXCR4 and corresponding mapping of sender and receiver cell states. **f,** Xenium data for the same-FFPE block (a consecutive section was unavailable), capturing homologous histopathological structures (top left). (top right) labelling of Xenium cell types based on label transfer from Wu et al. 2021. (bottom) Visualization of CXCL12 and CXCR4, confirming expression of these genes in Visium homologous areas.
