## Extended Data Fig. 9 for "Highly resolved tumor architecture via matched spatial and nucleus transcriptomics from a single tissue section"

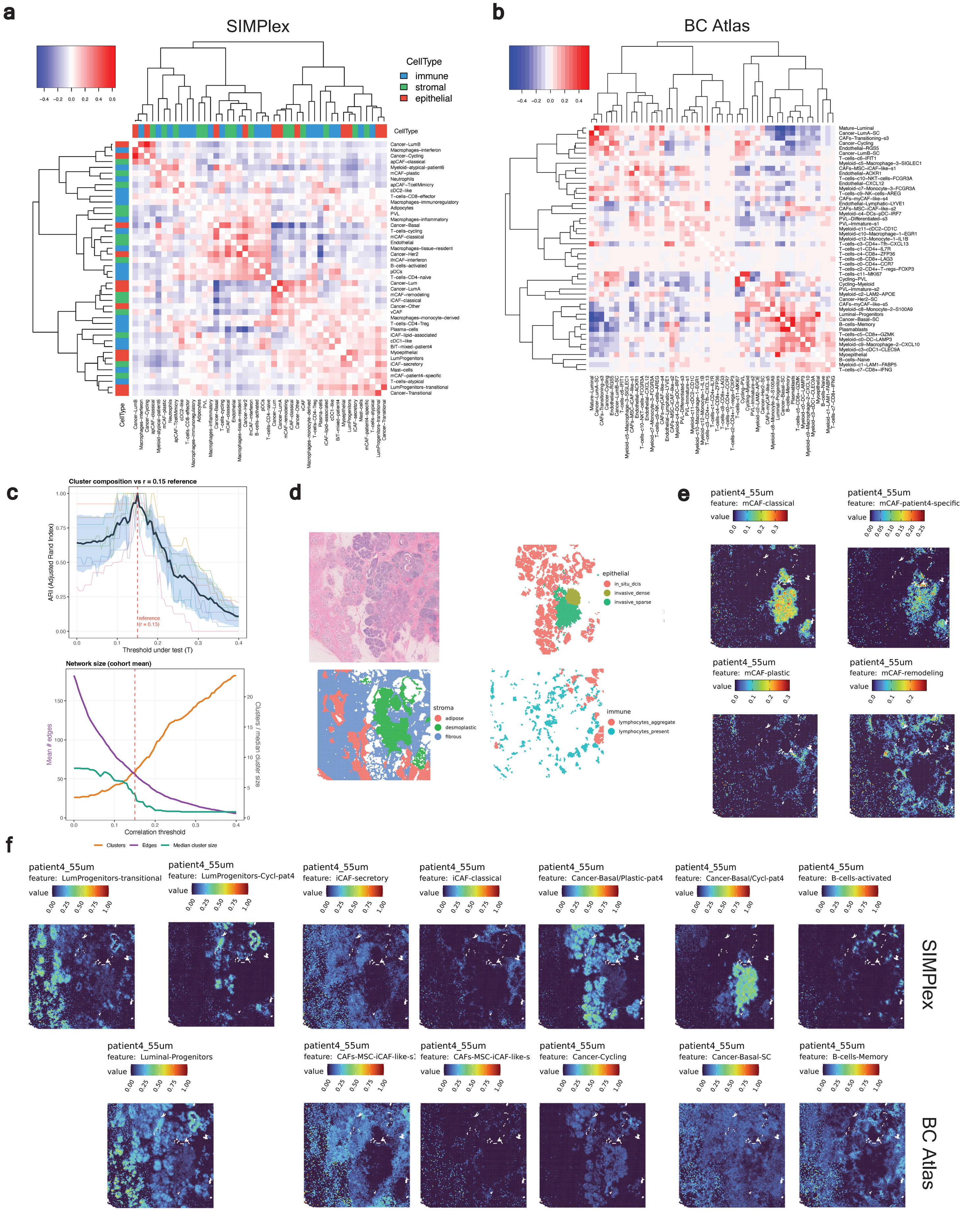


**Extended Data Fig.9 | Comparison of fine-grained cell state mapping. a,** Spatial colocalization heatmap of SIMPlex-mapped cell states summarised across all standard Visium sections. The scale bar denotes per-spot Spearman correlation coefficients. Patient-specific but biologically overlapping states were merged for interpretability (e.g., cancer-basal from Patients 4 and 5). **b,** Spatial colocalization heatmap of BC atlas-mapped cell states across all standard Visium sections. The scale bar denotes per-spot Spearman correlation coefficients. **c,** Sensitivity analysis of Spearman co-localization threshold. Reference partition: r = 0.15 (red dashed line; used in main niche analysis). Top: adjust-ed Rand index (ARI) between the Louvain partition at each T and the reference at r = 0.15 (faint lines, per sample; black line and blue ribbon, cohort mean ± s.d.; ARI = 1 at T = 0.15 by definition). Bottom: cohort-mean network size vs T - number of edges (purple, left y-axis); number of Louvain clusters and median cluster size (orange and teal, right y-axis; scaled for display). **d,** Original spot-level histopathology annotation for Patient 4 standard Visium data. **e,** Deconvolution maps of mCAF cell states in Patient 4. **f,** Comparison of deconvolution patterns between fine-grained SIMPlex cell states and the BC reference atlas. At this resolution, annotation nomenclature differs substantially and direct alignment is limited. A subset of comparable states was selected to illustrate differences in spatial segregation.
