## Extended Data Fig. 8 for "Highly resolved tumor architecture via matched spatial and nucleus transcriptomics from a single tissue section"

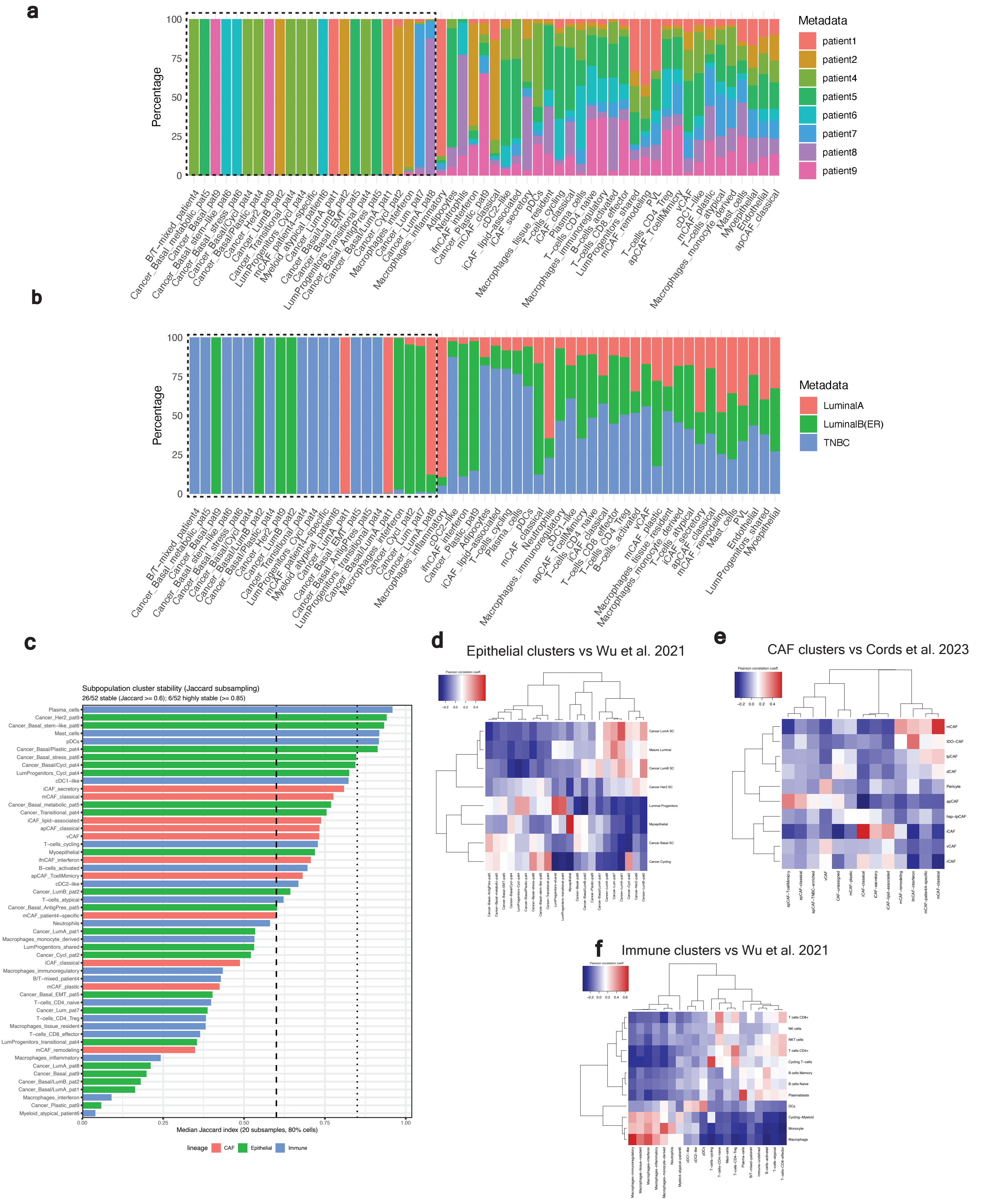


**Extended Data Fig.8 | Annotation of fine-grained SIMPlex cell states. a, b,** Patient and BC molecular subtype distribution across SIMPlex fine-grained cell states after subclustering an annotation of epithelial, immune, and CAF lineages. Dashed boxes denote cell states with exclusive or high degrees of patient-specificity. **c,** Cluster stability measured via median Jaccard similarity between each annotated subpopulation and its best-matching cluster after 20 rounds of 80% nuclei subsampling and lineage-specific re-clustering. Dashed and dotted horizontal lines indicate stability thresholds of 0.6 and 0.85, respectively. Colours denote lineage. **d,e,f,** Comparison of each annotated cell state with a relevant published atlas. Wu et al. 2021 was used for Epithelial and Immune populations and Cords et al. 2023 for CAF annotation. Cluster identities were compared by computing Pearson Correlations between average expression profiles across shared variable genes.
