## Extended Data Fig. 7 for "Highly resolved tumor architecture via matched spatial and nucleus transcriptomics from a single tissue section"

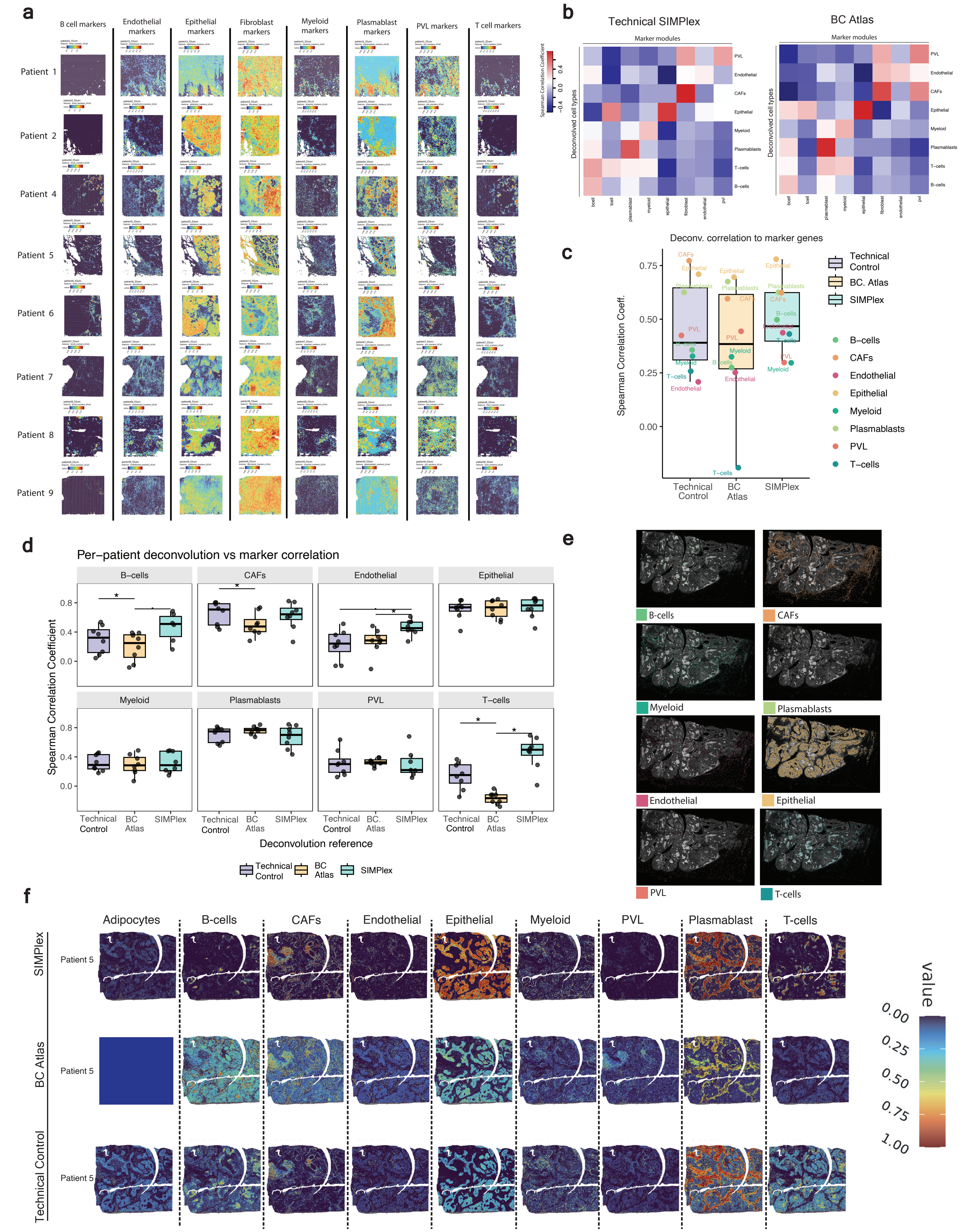


**Extended Data Fig.7 | a,** Spatial visualization of UCell marker gene scores in all standard Visium sections. **b,** Heatmap of per-spot SCC between marker gene scores and spatial deconvolution using Pooled SIMPlex reference (excluding corresponding-patient SIMPlex data from each Visium mapping) and reference BC atlas. **c,** Summary of SCCs between marker gene score and deconvolution with different references. Points represent whole-cohort SCCs for each cell type and matching cell type label. **d,** Per-cell type SCCs between marker gene score and deconvolution with different references. Points represent values for each section mapping (Wilcoxon signed-rank test and Benjamini-Hochberg correction; *: FDR < 0.05; (dot): FDR < 0.1). **e,** Spatial visualization of cell types in a consecutive Xenium section for patient 5. Cell types were identified via reference BC atlas label transfer. **f,** Spatial mapping of major cell types with different references in Patient 5 Visium HD data at 8μm bin resolution.
