## Extended Data Fig. 6 for "Highly resolved tumor architecture via matched spatial and nucleus transcriptomics from a single tissue section"

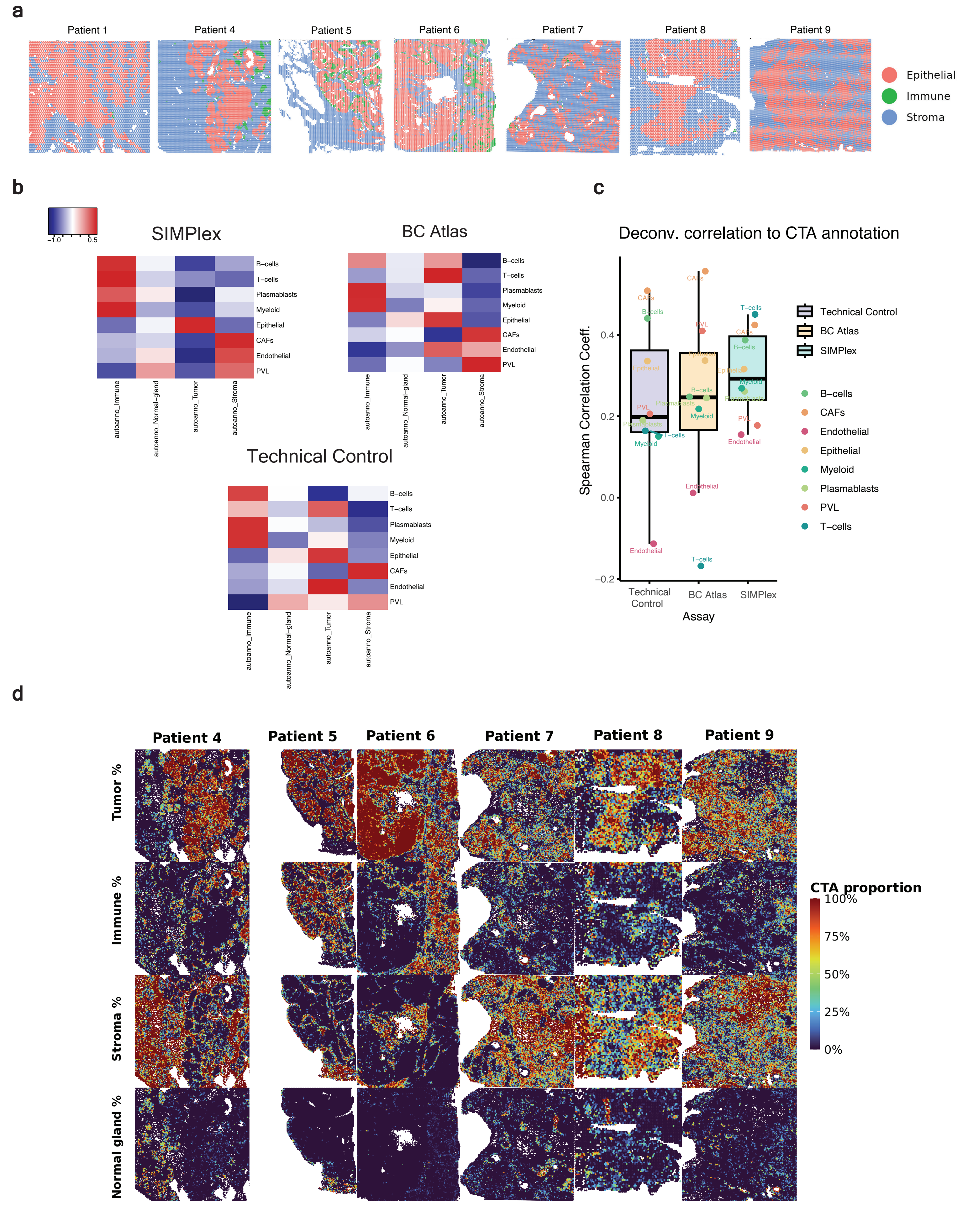


**Extended Data Fig.6 | Comparison of mapping to histopathology and computational tissue annotation. a,** Simplified spot-level histopathology annotation for standard Visium sections. **b,** Whole-cohort Spearman Correlation Coefficient (SCC) heatmap of mapping with different references and per-spot CTA cell counts. **c,** Summary of SCCs between CTA cell counts and deconvolution with different references. Points represent whole-cohort SCCs for each cell type and matching CTA label. **d,** Spatial mapping of CTA for the 6 patients analysed. Scale bar indicates cell count proportions.
