## Extended Data Fig. 5 for "Highly resolved tumor architecture via matched spatial and nucleus transcriptomics from a single tissue section"

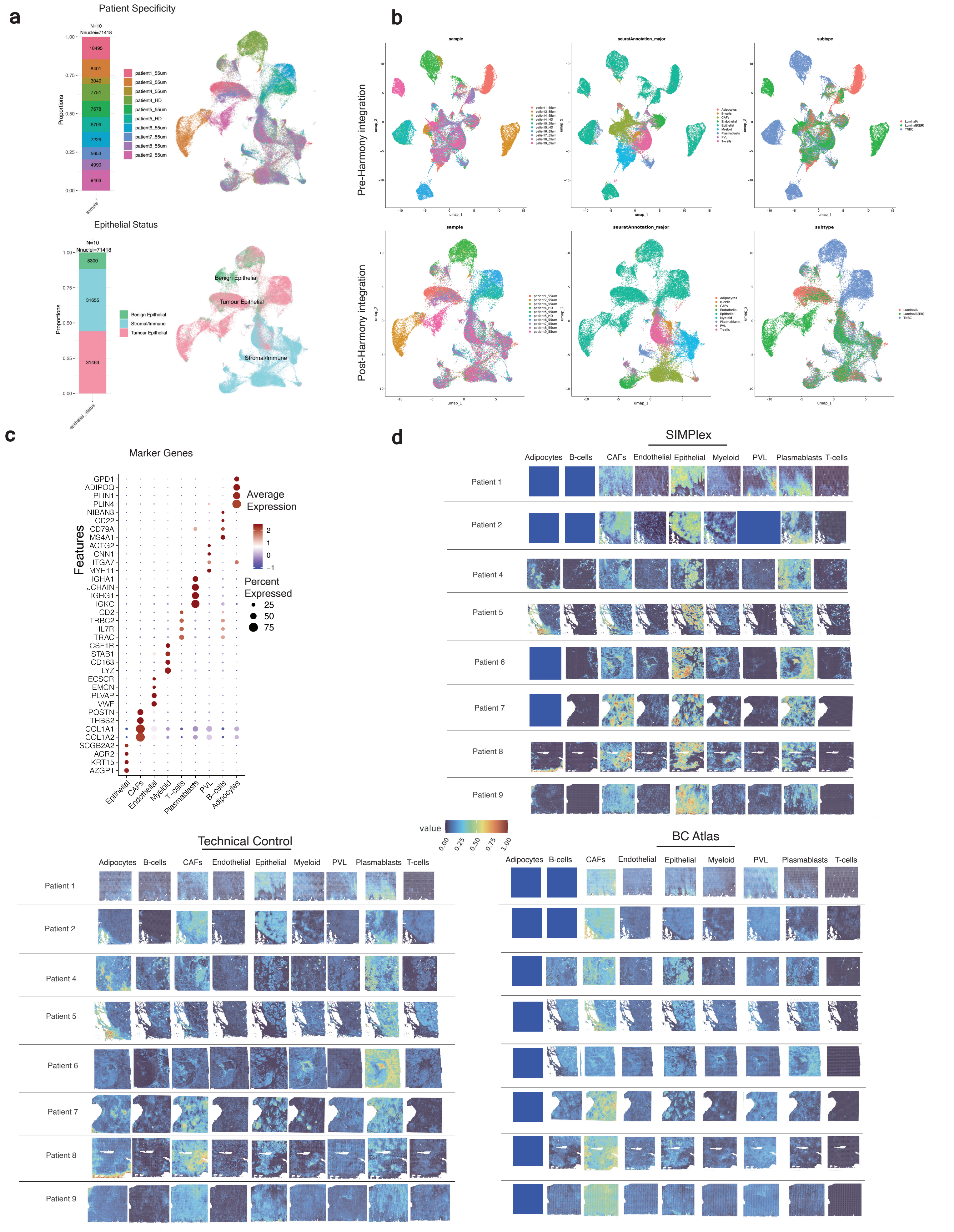


**Extended Data Fig.5 | Integrative analysis of cross-patient SIMPlex FFPE-BC data and spatial mapping of major cell types. a,** UMAP visualization of integrated SIMPlex BC dataset including data from all standard and HD Visium sections. Nuclei are colored by patient of origin (top) and epithelial status (tumour/benign) based on label transfer from the reference BC atlas (bottom). **b,** UMAP embeddings pre- and post-integration with Harmony. Nuclei are labelled by section of origin, annotation based on BC Atlas label transfer, and molecular subtype. **c,** Dot-plot showing marker genes confirming SIMPlex cell type annotations. Dot size represents the percentage of nuclei expressing each gene, and color indicates average expression. **d,** Spatial mapping of major cell types using: (right) patient-matched SIMPlex data exclusively; (bottom) BC Atlas reference; (left) Technical Control.
