## Extended Data Fig. 4 for "Highly resolved tumor architecture via matched spatial and nucleus transcriptomics from a single tissue section"

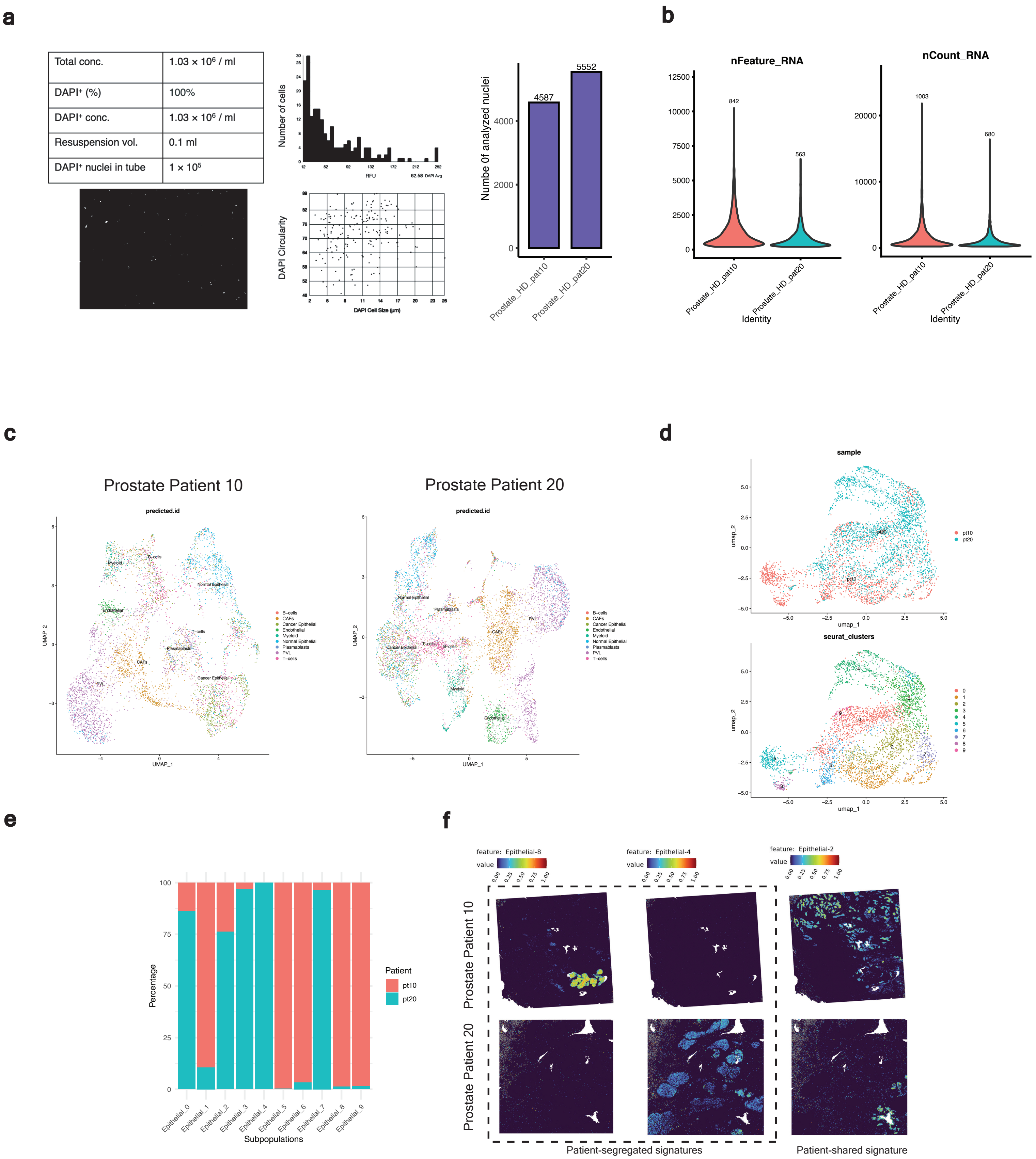


**Extended Data Fig.4 | SIMPlex Visium HD profiling of FFPE prostate cancer tissue from two patients.**
**a,** SIMPlex workflow applied to two 5 µm FFPE-prostate sections, barcoded separately and pooled prior to nuclei isolation.Countess nuclei counting output following DAPI-based sorting and number of nalazyed nuclei after data pre-processing. **b,** SIMPlex-nuclei metrics displaying unique genes (nFeature_RNA) and UMI counts (nCount_RNA). Mean values per patient are presented on top of each violin. **c,** UMAP visualization of prostate SIMPlex data for each patient/ section. **d,** Integration and subclustering of epithelial lineage of the two SIMPlex prostate sections. Nuclei are labelled by patient of origin (top) and cluster (bottom). **e,** Sample distribution of epithelial clusters shown in (d). **f,** Spatial deconvolution of integrated epithelial clusters onto Visium HD data (8µm resolution bin). Illustrative clusters to show patient-specificity and shared patterns are displayed.
