## Extended Data Fig. 3 for "Highly resolved tumor architecture via matched spatial and nucleus transcriptomics from a single tissue section"

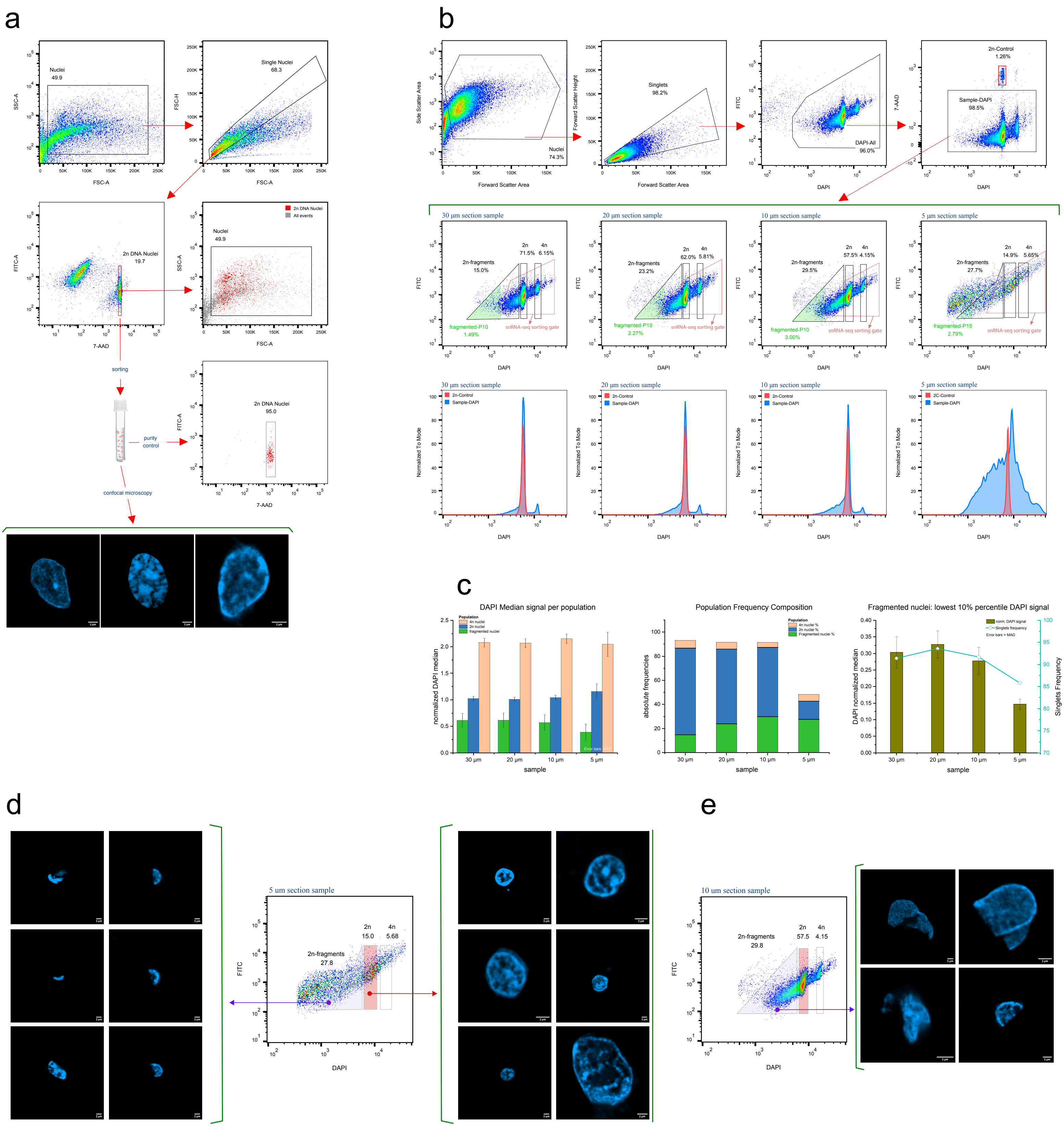

**Extended Data Fig. 3 | Nuclei integrity assessment across section thicknesses.**
Nuclei integrity was assessed in 5, 10, 20, and 30 µm-thick sections of FFPE breast cancer (BC) tissue. **a,** Gating strategy for the control sample. Nuclei were gated sequentially: Side Scatter Area vs. Forward Scatter Area to identify nuclei, Forward Scatter Height vs. Forward Scatter Area to select singlets, and 7-AAD staining to identify 2n intact nuclei. The sorted 2n control nuclei were assessed for purity by FANS and confocal microscopy prior to spiking-in to samples. **b,** FANS analysis of the four samples (5, 10, 20, 30 µm), each spiked with the pre-sorted 2n control nuclei and stained with DAPI. Top row: gating strategy for single nuclei population selection and separation of control from sample nuclei. Middle row: segmentation of sample nuclei (excluding the control nuclei) into fragmented, 2n, and 4n populations. The red gates denote the snRNA seq sorting gate including only the larger fragmented nuclei, 2n and 4n. Bottom row: DAPI-channel histograms for each of the four samples (blue), with the 2n control nuclei signal (red) overlay. **c,** Statistical analysis of flow cytometric nuclei integrity data. Left: normalized median DAPI signal for the three nuclei segments (fragmented, 2n, 4n) across all four samples, normalized to the median DAPI signal of the 2n control nuclei. Middle: absolute frequencies of the three nuclei populations to their parent gate Sample-DAPI across the four samples. Right: normalized median DAPI signal of the lowest 10th percentile of fragmented nuclei (events under green area in middle row, panel b), plotted alongside Singlets frequencies (light blue line, right Y-axis). **d,** Confocal images of sorted nuclei from the 5 µm section sample: fragmented nuclei (left) and 2n nuclei (right). **e,** Confocal images of sorted fragmented nuclei from the 10 um section sample.
