## Extended Data Fig. 2 for "Highly resolved tumor architecture via matched spatial and nucleus transcriptomics from a single tissue section"

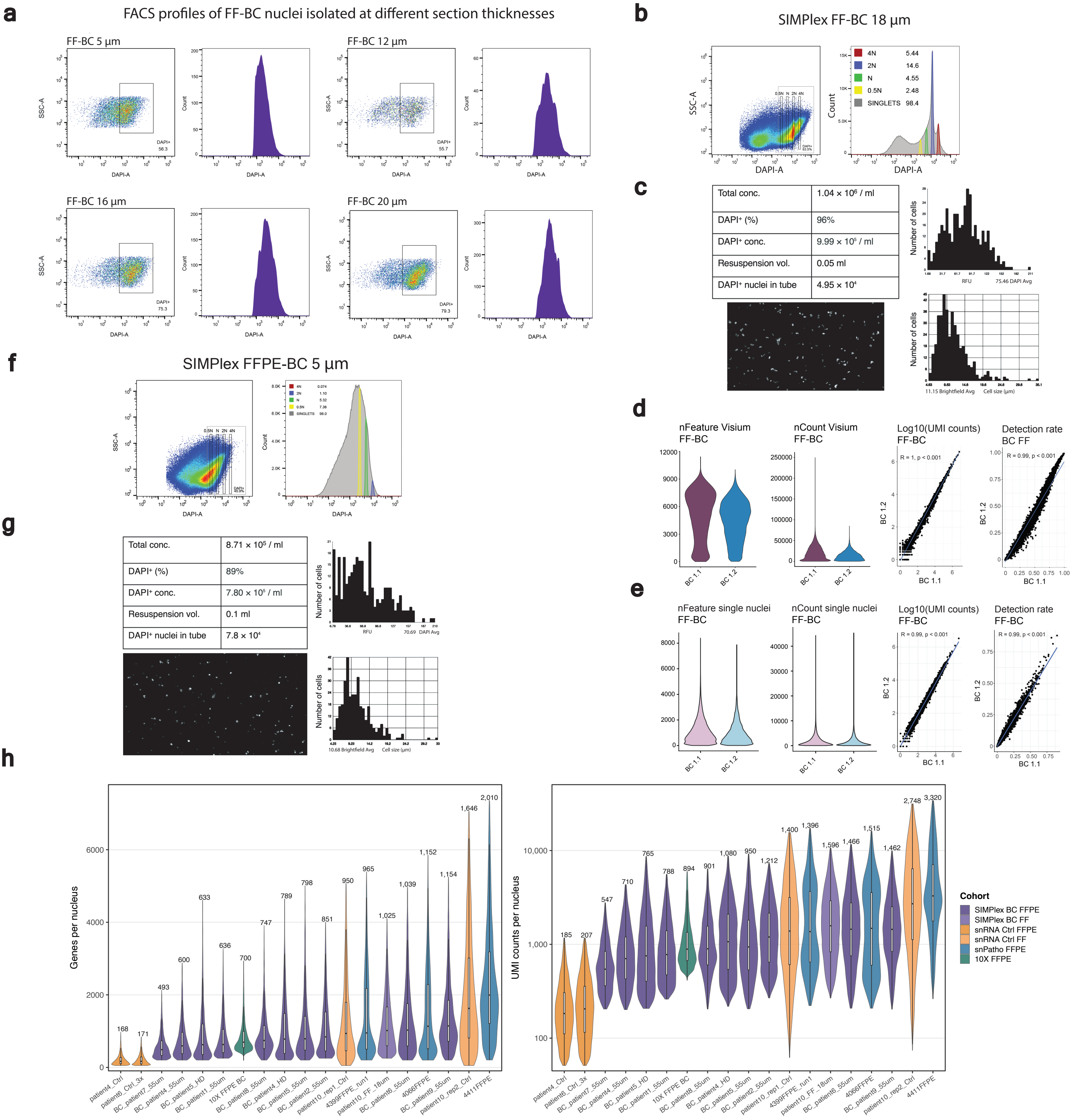


**Extended Data Fig.2 | Nuclei and transcriptomic QC for SIMPlex across FF and FFPE breast cancer tissue. a,** FACS profiles of nuclei isolated from FF-BC tissue sections of varying thicknesses, used to assess nuclei quality prior to the full SIMPlex workflow. Side-scatter versus DAPI plots and the corresponding DAPI histograms are shown for each thickness. All data are displayed after nuclei gating in side-scatter versus forward-scatter space with singlet selection applied. DAPI⁺ values indicate the proportion of events within the DAPI⁺ gate. **b,c,d,e,** Full SIMPlex workflow applied to two 18 µm FF-BC sections, barcoded separately and pooled prior to nuclei isolation. (b) Side-scatter versus DAPI profiles and corresponding DAPI histogram after nuclei and singlet gating. (c) Representative Countess nuclei counting output following DAPI-based sorting. (d) Visium quality metrics from the matched spatial dataset, including distributions of detected genes (nFeature) and molecules (nCount) per spot for BC 1.1 and BC 1.2, as well as gene-gene UMI correlations ($log_{10}$-transformed) and detection-rate comparisons between the two sections. (e) Corresponding single-nucleus RNA-seq metrics for the same sections, including nFeature and nCount distributions per nucleus, gene-gene UMI correlations, and detection-rate comparisons. **f,g,** Represenative SIMPlex FFPE-BC 5 µm QC from two sections, barcoded separately and pooled prior to nuclei isolation.(f) Side-scatter versus DAPI profiles and corresponding DAPI histogram after nuclei and singlet gating. (g) Representative Countess nuclei counting output following DAPI-based sorting. **h,** Distributions of detected genes (nFeature_RNA) and UMI counts (nCount_RNA) per nucleus across SIMPlex samples (standard Visium and HD), compared with public FFPE snRNA-seq datasets (10x FFPE breast cancer and snPATHO-FFPE). Controls (yellow) reflect samples that did not undergo Visium before nuclei isolation from a tissue section. Median values for each dataset are indicated above the violins. Data has been clipped at the 99th percentile for visibility.
