## Extended Data Fig. 1 for "Highly resolved tumor architecture via matched spatial and nucleus transcriptomics from a single tissue section"

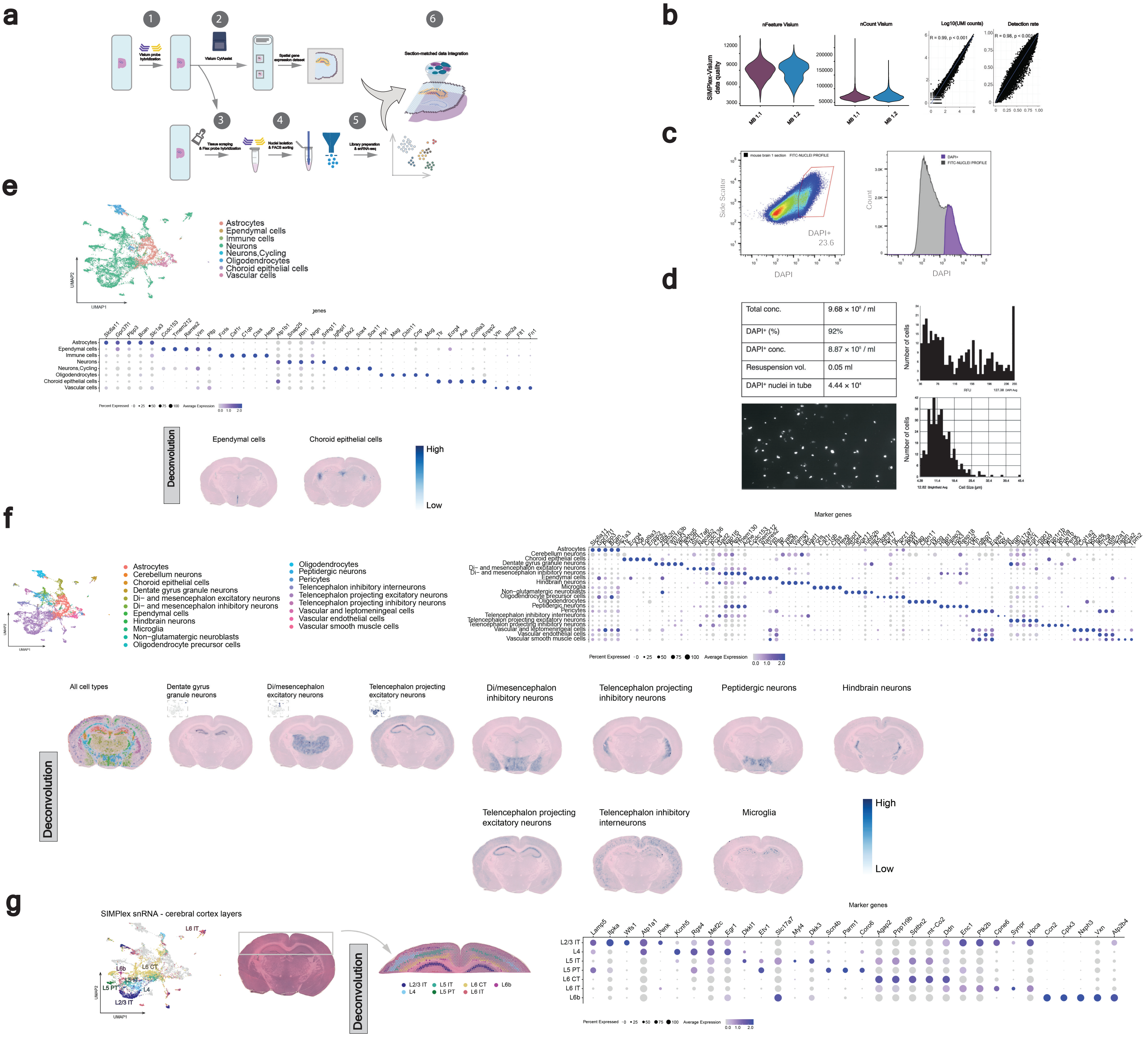

**Extended Data Fig.1 I SIMPlex workflow and validation in mouse brain.**

**a,** (1) An H&E-stained tissue section prepared from fresh-frozen or FFPE blocks mounted on a Superfrost Plus slide is hybridized with Visium FFPE probes. (2) The slide is processed on the Visium CytAssist instrument, which transfers probe-bound transcripts to the Visium capture slide for spatial library preparation and generation of the spatial gene expression dataset. (3) The original H&E section remaining on the Superfrost slide is scraped into LoBind tubes and Flex whole-transcriptome probes are added for hybridization in suspension. (4) Lysis and filtration release nuclei, followed by DAPI staining and FACS sorting of nuclei. (5) Sorted nuclei undergo Flex-based library preparation and sequencing to generate single-nucleus transcriptomes. (6) Spatial and single-nucleus transcriptomes derived from the same physical tissue section are integrated to enable matched-section deconvolution and multimodal analysis. **b,c,d,** SIMPlex workflow applied to two 12 µm FF-mouse brain sections, barcoded separately and pooled prior to nuclei isolation. **b,** Matched-Visium QC metrics for MB 1.1 and MB 1.2, including distributions of detected genes (nFeature) and molecules (nCount) per spot, gene–gene UMI count correlations ($log_{10}$-transformed), and gene detection-rate comparison. Detection rate is defined as the proportion of spots with detected UMIs. Pearson correlation coefficients and two-sided P values are shown; MB 1.1 (mouse brain section 1), MB 1.2 (mouse brain consecutive section 2). **c,** Representative flow cytometry Side Scatter versus DAPI profile and corresponding DAPI histogram. Data are plotted after hierarchical gating was applied for Nuclei (SSC vs FCS), Singlets (FSC-A vs FSC-H) and FITC Filtering (SSC vs FITC) for excluding debris and high autofluorescence outliers. The DAPI⁺ value indicates the proportion of events within the DAPI⁺ gate.(**d**) Representative Countess nuclei counting output following DAPI-based sorting. **e,** UMAP visualization of major cell types identified in the SIMPlex-snRNA mouse brain dataset. Cell types are annotated by label transfer from the Mouse Brain Atlas. A dot plot shows marker gene expression across these cell types, with dot size indicating the percentage of expressing cells and color intensity reflecting average expression. Matched Visium sections deconvolved with SIMPlex-snRNA references are shown. **f,** As in panel e, but showing more fine-grained SIMPlex-snRNA cell-type annotations and their mapped distribution across matched Visium sections. **g,** UMAP of SIMPlex-snRNA nuclei annotated (using the Allen Mouse Brain Atlas, cortex-focused), with cortical layer subtypes (L2/3 IT, L4, L5 IT, L5 PT, L6 CT, L6 IT, L6b), alongside deconvolution of the matched Visium section to reveal laminar architecture. Marker gene expression is shown as a dot plot, with dot size indicating percentage expression and color reflecting average expression.
